# Ribosomal Frameshifting Selectively Modulates the Assembly, Function, and Pharmacological Rescue of a Misfolded CFTR Variant

**DOI:** 10.1101/2023.05.02.539166

**Authors:** Patrick Carmody, Francis J. Roushar, Austin Tedman, Wei Wang, Madeline Herwig, Minsoo Kim, Eli F. McDonald, Karen Noguera, Jennifer Wong-Roushar, Jon-Luc Poirier, Nathan B. Zelt, Ben T. Pockrass, Andrew G. McKee, Charles P. Kuntz, S. Vamsee Raju, Lars Plate, Wesley D. Penn, Jonathan P. Schlebach

## Abstract

The cotranslational misfolding of the cystic fibrosis transmembrane conductance regulator chloride channel (CFTR) plays a central role in the molecular basis of cystic fibrosis (CF). The misfolding of the most common CF variant (ΔF508) remodels both the translational regulation and quality control of CFTR. Nevertheless, it is unclear how the misassembly of the nascent polypeptide may directly influence the activity of the translation machinery. In this work, we identify a structural motif within the CFTR transcript that stimulates efficient -1 ribosomal frameshifting and triggers the premature termination of translation. Though this motif does not appear to impact the interactome of wild-type CFTR, silent mutations that disrupt this RNA structure alter the association of nascent ΔF508 CFTR with numerous translation and quality control proteins. Moreover, disrupting this RNA structure enhances the functional gating of the ΔF508 CFTR channel at the plasma membrane and its pharmacological rescue by the CFTR modulators contained in the CF drug Trikafta. The effects of the RNA structure on ΔF508 CFTR appear to be attenuated in the absence of the ER membrane protein complex (EMC), which was previously found to modulate ribosome collisions during “preemptive quality control” of a misfolded CFTR homolog. Together, our results reveal that ribosomal frameshifting selectively modulates the assembly, function, and pharmacological rescue of a misfolded CFTR variant. These findings suggest interactions between the nascent chain, quality control machinery, and ribosome may dynamically modulate ribosomal frameshifting in order to tune the processivity of translation in response to cotranslational misfolding.

**Significance:** Many diseases stem from imbalances between protein synthesis and degradation that arise from mutations and/ or cellular stressors. The molecular mechanisms responsible for such lapses in cellular proteostasis often coincide with aberrant regulation of protein translation. Here, we identify a structure within the transcript encoding the CFTR chloride channel that allows the ribosome to halt translation in response to its cotranslational misfolding. We show that this motif modifies the assembly, function, and pharmacological properties of the most common cystic fibrosis variant. This crosstalk between the ribosome and nascent polypeptide allows the ribosome to adjust its activity to prevent the synthesis of misfolded proteins. These findings suggest ribosomal frameshifting and premature translational termination plays a fundamental role in protein quality control.

## Introduction

Cystic fibrosis (CF) is a disease of protein homeostasis (proteostasis) that arises from the defective biosynthesis, folding, and/ or function of a chloride channel known as the cystic fibrosis transmembrane conductance regulator (CFTR).^1,2^ The most common CF mutation (ΔF508) induces CFTR misfolding by decoupling the folding of its subdomains during translation.^3,4^ This cotranslational misassembly reaction enhances the retention and degradation of the CFTR protein within the endoplasmic reticulum (ER) and ultimately reduces the trafficking of the functional protein to the plasma membrane. This lapse in protein quality control (QC) coincides with the remodeling of CFTR proteostasis network and changes in the dynamics of CFTR translation.^5-11^ Nevertheless, the precise chain of events involved in the crosstalk between the conformational state of nascent CFTR, the activity of the translation machinery, and the interaction of the nascent chain with various components of the cellular proteostasis network is not fully understood.

CFTR QC begins during its cotranslational folding at the ribosome-translocon complex.^12^ Vectorial folding of its five subdomains is orchestrated, in part, by positional variations in translation kinetics that are coordinated by rare codons,^9^ the relative abundance of certain translation factors,^5,7^ and the presence of certain mRNA secondary structures.^13,14^ Together, these effectors help the ribosome tailor its activity to the kinetic constraints of cotranslational CFTR folding. Indeed, the effects of CF mutations on CFTR folding and assembly are highly sensitive to changes in translational dynamics. For instance, suppressing translation initiation reduces the density of ribosomes on the ΔF508 transcript in a manner that partially rescues its stability, cellular trafficking, and function.^7,11^ The cotranslational misfolding of a ΔF508 CFTR homolog also appears to modulate translation in *cis* by enhancing ribosome collisions and triggering “preemptive quality control”-a cotranslational QC pathway mediated by the ER membrane protein complex (EMC).^11^ Though the later finding suggests the intrinsic activity of the ribosome is sensitive to conformational transitions in the nascent chain, it remains unclear whether cotranslational misfolding events are capable of directly altering translation in real time.

We recently found that cotranslational folding mechanically modulates a translational recoding mechanism known as -1 programmed ribosomal frameshifting (−1PRF) during viral polyprotein biosynthesis.^15,16^ These findings reveal that, under certain circumstances, the ribosome exhibits an enhanced propensity to slip into alternative reading frames in response to conformational transitions in the nascent polypeptide chain.^17^ In the following, we evaluate whether similar feedback occurs during CFTR biosynthesis. We first identify a structured segment within the region of the CFTR transcript that stimulates ribosomal frameshifting (RF). We then show that the disruption of this motif selectively remodels the interactome of ΔF508 CFTR in a manner that partially restores its expression and function. These modifications, which have minimal impact on WT CFTR, also enhance the pharmacological rescue of ΔF508 CFTR by the leading CF therapeutic Trikafta. Our results show that this feedback hinges on the EMC, suggesting this mechanism is involved in preemptive QC. Our results suggest this RF site acts a QC-mediated translational “kill switch” that selectively promotes the premature termination of translation in response to the cotranslational misfolding of the nascent chain.

## Results

### Discovery of an Active -1 Ribosomal Frameshift Site in the CFTR Transcript

Our recent findings show that the mechanical forces generated by cotranslational folding can stimulate -1PRF,^15,16^ which minimally requires a slippery sequence within the mRNA and an adjacent stem-loop.^18^ To identify potential RF sites in the CFTR transcript we first searched for consensus or near-consensus X_1_ XXY_4_ YYZ_7_ slippery sequences, where XXX and YYY are nucleobase triplets and Z can be any nucleobase.^19-21^ We identified five consensus X_1_ XXY_4_ YYZ_7_ sites and eight other near-consensus sites distributed throughout the transcript (Figs. 1A & S1). Three of these sites are clustered within the region encoding nucleotide binding domain 2 (NBD2) and are proximal to two predicted RNA stem-loops (Fig. 1A). A uridine rich sequence referred to here as slip-site A (UUU AUU UUU UCU, SSA) lies upstream of the canonical slip-site B (A_1_ AAA_4_ AAC_7_, SSB). SSB is eight nucleotides upstream of a small, predicted stem-loop (SL2). We also identified a third putative non-canonical slip site (A_1_ GAA_4_ AUA_7_, SSC) downstream of SSB that is positioned seven nucleotides upstream of a much longer putative stem-loop (SL2); an ideal spacing for the stimulation of -1PRF.^18^

**Figure 1.**
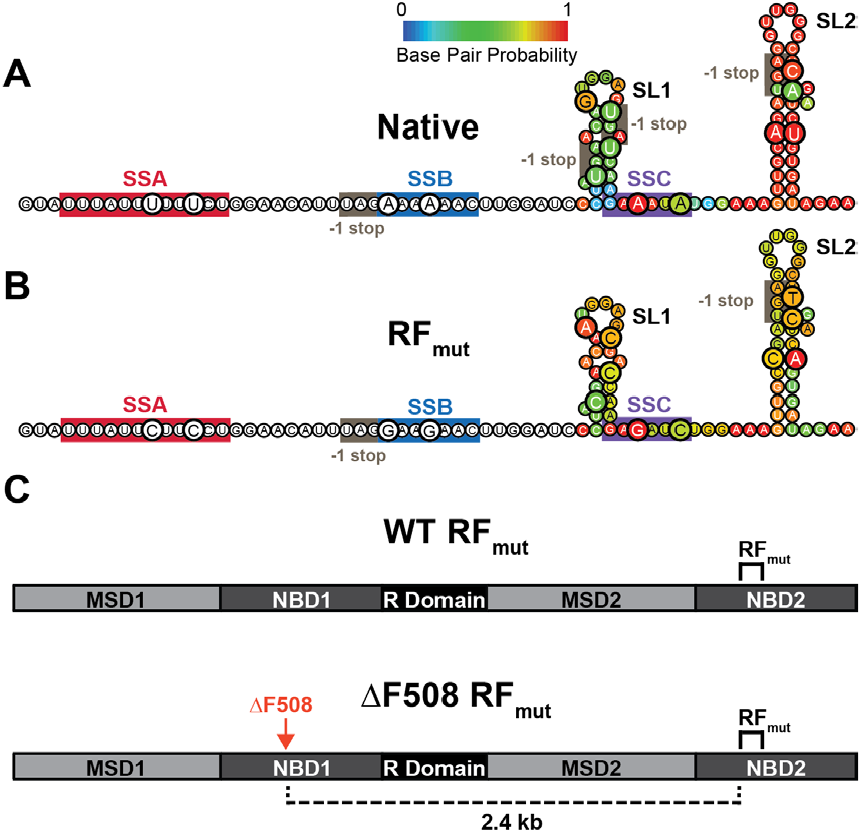
Putative mRNA structures within the CFTR ribosomal frameshift motif. A) The predicted secondary structure downstream of SSB is shown. Bases are colored according to their predicted base pair probabilities as determined by Vienna RNAfold. B) Secondary structure predictions for a construct bearing 14 silent mutations (RF_mut_) are shown for the sake of comparison. Mutated bases are enlarged and bases are colored according to their predicted base pair probabilities as determined by Vienna RNAfold in both panels A and B, for reference. The positions of the three putative slip sites and the downstream stop codons in the -1 reading frame are indicated. C) A schematic depicts the layout of the full-length WT and ΔF508 CFTR transcripts bearing the RF_mut_ modifications.

Given that ribosome collisions can occur during CFTR translation^11^ and that they are known to modulate -1PRF,^22^ we hypothesized that RF could occur at one or more of these sites. To determine whether efficient -1 RF occurs at any of these positions, we generated a series of bicistronic reporters in which a short segment of the CFTR transcript bearing these features is inserted between an upstream Renilla luciferase (rLuc, expression control) and a downstream -1 firefly luciferase (fLuc, -1 RF reporter, Fig. 2A).^23^ We generated three versions of this reporter that produce fLuc in response to frameshifting at any of these slip-sites (SSABC), or conditionally at B or C (SSBC), or at C only (SSC) by knocking out various -1 frame stop codons downstream of these slip-sites (Fig. 2A orange). Transient expression of the SSABC reporter in a human bronchial epithelial cell line (CFBE41o-) generates a detectable fLuc frameshift signal 4-fold over the no-insert control baseline. rLuc intensities are similar for all three reporters, suggesting all three reporters exhibit similar expression under these conditions (Fig. 2B). fLuc intensities are also quite similar for all three reporters (Fig. 2C), which implies most frameshifting occurs at SSC. Consistent with this interpretation, we find that the fLuc signal is ablated by mutations that disrupt SSC (SSC_mut_, Fig. 2C). We note that we ruled out splicing artifacts^24^ by RT-PCR and found that the frameshift signal can also be partially suppressed by stop codons in either upstream of SSC in the 0-frame (5′Ter) or downstream of SSC in the -1 frame (3′Ter, Figs. 2D & S2), which confirms that these signals arise from a genuine RF. By normalizing the rLuc and fLuc signals, we estimate ribosomal frameshifting occurs with an efficiency of 2.9 ± 0.2% at SSC, which is comparable to levels achieved during translation of the canonical HIV gag-pol motif (5.2 ± 0.4%, Fig. 2D). Similar frameshifting also occurs in HEK293T cells (Fig. S3). Together, these results identify an active -1 RF motif within the human CFTR transcript.

**Figure 2.**
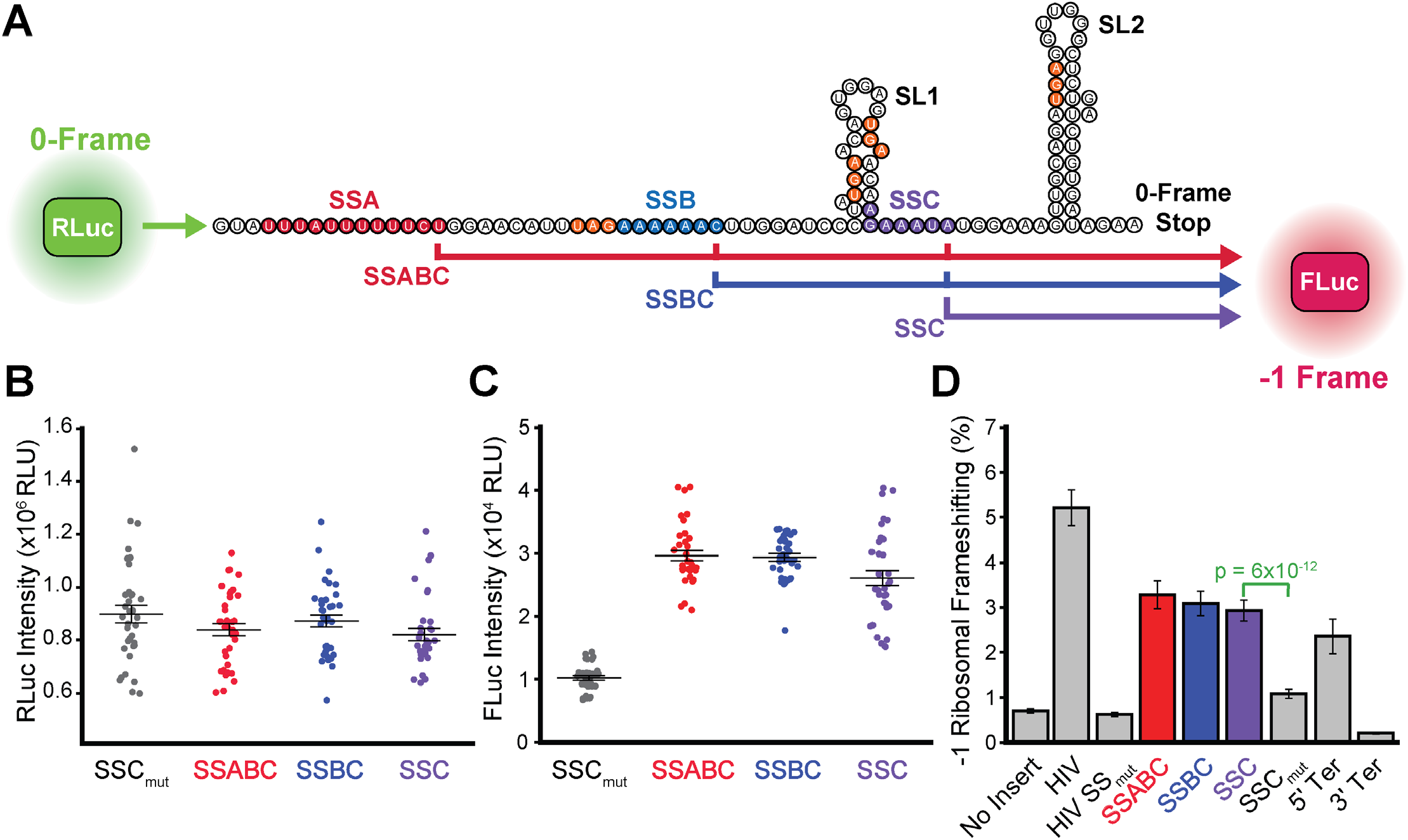
Stimulation of ribosomal frameshifting by structural elements within the CFTR transcript. The efficiency of ribosomal frameshifting during the translation of a structured region within the CFTR transcript was assessed in CFBE41o-cells using a bicistronic dual luciferase reporter system. (A) Cartoon depicts the logic of a series of reporter constructs that generate Renilla Luciferase (RLuc) upon translation initiation and a firefly Luciferase (FLuc) in response to a -1 ribosomal frameshift. The positions of the potential slip-sites A (red), B (blue), and C (purple) are shown. Stop codons in the -1 frame are indicated in orange. Putative secondary structures generated by Vienna RNAfold for the 75 bases beginning 5 bases downstream of SSB are shown for reference. (B, C) Dot plots indicate the raw RLuc and FLuc luminescence intensities within the lysates of CFBE41o-cells transiently expressing these constructs, respectively. Results were taken from three biological replicates with 4 transfections per sample each. Central hashes represent the average intensities and whiskers reflect their standard deviation. (D) A bar graph depicts the average -1 ribosomal frameshifting efficiencies for each construct in CFBE41o-cells. Error bars reflect the standard deviation to serve as a measure of precision.

### Impact of Ribosomal Frameshifting on the CFTR Interactome

Unlike viral -1PRF events that generate functionally distinct proteins, a -1 frameshift at SSC should only fuse five non-native residues onto nascent CFTR prior to translational termination and truncation in the middle of NBD2 (Fig. 1A). Based on this consideration, we hypothesized that the RF motif serves to promote frameshifting and the premature termination of translation in response to cotranslational CFTR misassembly. To determine whether this motif is involved in assembly, we assessed how this RNA structure influences the interactomes of WT and ΔF508 CFTR. Briefly, we designed a series of mutations to disrupt the slip-sites and weaken the stem-loops within NBD2 while preserving the amino acid sequence of CFTR in the 0-frame (RF_mut_, Fig. 1B). We then incorporated these modifications into the full length CFTR (Fig. 1C) and used affinity purification-mass spectrometry (AP-MS) based interactome profiling to determine the effects of these silent modifications on the interactions formed by WT and ΔF508 CFTR.^5,25,26^ In agreement with previous reports,^5,25^ the ΔF508 mutation significantly enhances the interaction of CFTR with numerous proteostasis factors (Fig. 3A & Table S1). By comparison, the silent mutations in NBD2 have minimal impact on the WT CFTR interactome (WT-RF_mut_, Fig 3A). In contrast, these same mutations significantly alter the propensity of ΔF508 CFTR to interact with several translation factors and quality control proteins (ΔF508-RF_mut_, Fig. 3A). Consistent with our hypothesis, the disparate effects of this transcript modification on WT and ΔF508 suggests the native RNA structure in NBD2 selectively alters CFTR translation and assembly in response to misfolding. We note that the ΔF508 mutation is ∼2.4 kb upstream of the ribosomal frameshift site in NBD2 (Fig. 1C), which suggests these differences likely arise from the effects of the ΔF508 mutation on the nascent CFTR protein rather than perturbation of the mRNA structure.

**Figure 3.**
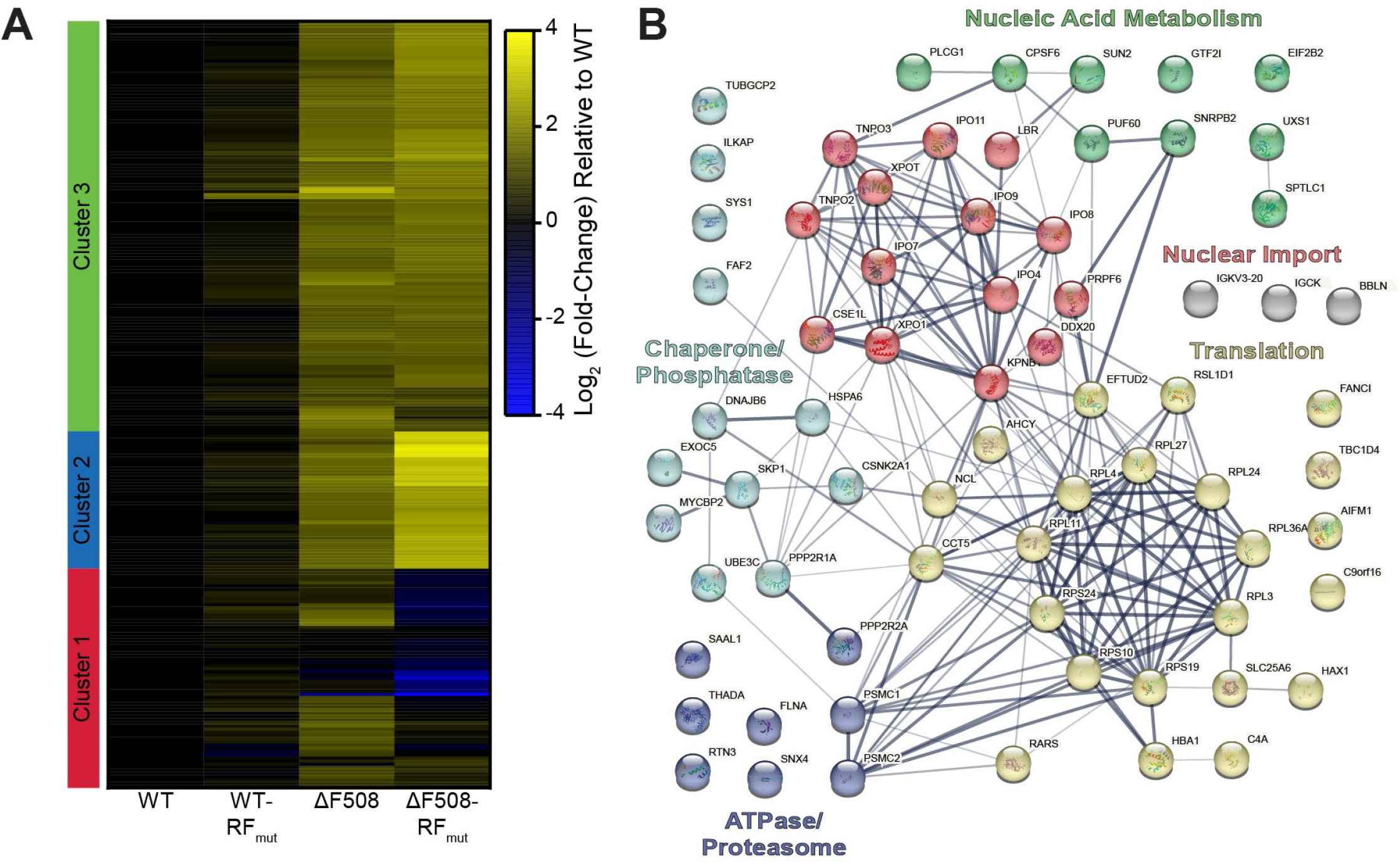
Impact of ribosomal frameshifting on the CFTR interactome. Affinity purification-mass spectrometry was employed to compare the effects of the silent RF_mut_ mutations on the interactome of WT and ΔF508 CFTR. A) A heatmap indicates the log_2_ (fold-change) in interactions of CFTR variants relative to WT. Increases in the association of interactors with CFTR variants relative to WT are indicated in yellow while blue indicates a decrease in relative abundance of interactors. Black indicates no change relative to WT. Scale shown in log_2_ fold change over WT abundance. Proteins are organized into 3 clusters according to a hierarchical clustering analysis that groups interactors based on changes in abundance. B) A network map depicts the relationships between interactors in cluster 1, which includes interactors the exhibit the largest changes in the context of ΔF508 RF_mut_ relative to ΔF508. Lines indicate known protein-protein interactions in the String database (human). Line widths indicate the strength of data support. The colors of nodes reflect the identity of sub-clusters from K-means clustering of the interactors within hierarchical cluster 1. The labels for each color summarize the most common classes of proteins within each sub-cluster. Interactors in gray were missing from the String database and were manually added in as isolated nodes.

There are several key differences in the interaction profiles of ΔF508 and ΔF508-RF_mut_ CFTR (Table S1). We first note that about half of the aberrant interactions formed by ΔF508 CFTR are unaffected by these silent mutations (see Cluster 1 in Fig. 3B & Clusters 2-3 in Fig. S4-S5). As both genetic constructs encode the same protein in the 0-frame, we suspect these common interactions may arise from the post-translational effects of the ΔF508 mutation on the CFTR protein. Nevertheless, ΔF508-RF_mut_ exhibits significantly attenuated associations with eight ribosomal proteins (Fig. 3B, Table S1). These silent mutations also appear to modulate QC, as ΔF508-RF_mut_ exhibits both weaker interactions with certain chaperones and co-chaperones (e.g. CANX, HspA6, BAG2, and DNAJB6) and stronger interactions with others (e.g. HspB1, HspA4, and DNAJA3 & Fig. S5). These QC modifications also coincide with a decrease in the interaction of ΔF508-RF_mut_ with the UBE3C ubiquitin ligase, the ER-PHAGY protein RTN3, and various components of the 19S proteasome complex (Table S1), which suggests the RF motif ultimately modifies CFTR degradation. We note these silent mutations enhance the interaction of ΔF508 CFTR with two components of the eIF3 complex (i.e. eIF3c and eIF3e, Cluster 3, Fig. S4, Table S1), which is known to modify the proteostatic effects of the ΔF508 mutation.^7^ Finally, these silent mutations also attenuate interaction with a subunit of the eIF2 complex (eiF2B2), whose alpha subunit was recently seen to regulate -1PRF in SARS-CoV-2. Overall, these results suggest the ribosomal frameshift site in NBD2 acts as a junction where cotranslational misfolding triggers changes in CFTR translation, QC, and degradation.

### Impact of Ribosomal Frameshifting on CFTR Expression

To determine how these changes in cotranslational assembly impact CFTR proteostasis, we compared the effects of the RF_mut_ modification on ΔF508 and WT CFTR expression. Western blot analyses reveal that modifications of the RF motif have minimal impact on the maturation of WT or ΔF508 CFTR in both CFBE41o- and HEK293T cells as judged by the relative abundance of the mature (C band) and immature (B band) glycoforms (Figs. 4 A-C, Fig. S6). This observation is perhaps unsurprising given that the RF_mut_ modification does not ultimately change the amino acid sequence of the dominant CFTR translation products, which are still subject to many of the same post-translational QC interactions (Fig. 3A). The RF_mut_ modification also appears to have little, if any, impact on total CFTR levels in CFBE41o-cells, as detected by western blotting (Fig. 4D). Any subtle differences in expression under these conditions are likely masked by the large variations in the transcript levels generated by transient transfections (Fig. S7). To quantitatively compare cellular CFTR accumulation, we used flow cytometry to compare single-cell surface and intracellular CFTR immunostaining levels in HEK293T cells (see *Methods*). Flow cytometry measurements confirm that disrupting the RF motif does has no impact on the WT CFTR expression and does not rescue the plasma membrane expression of ΔF508 CFTR (Fig. 4 E-F). However, the RF_mut_ modification does increase intracellular ΔF508 levels by 19 ± 11% (*p* = 0.007, Fig. 4 E-F). Notably, the intracellular immunostaining intensity of ΔF508-RF_mut_ is indistinguishable from that of WT (Fig. 4E), which suggests this motif normally contributes to the reduced accumulation of ΔF508 CFTR in the ER. While all trends in expression were generally consistent in CFBE41o- and HEK293T cells (Figs. 4 & S6), this enhanced intracellular accumulation of ΔF508-RF_mut_ CFTR was not observed in the context of CRISPR knockout cells lacking the EMC (Figs. S8 & S9). This observed connection to the EMC potentially suggests this feedback may play a role in preemptive QC. Though changes in expression are subtle, our results reveal that the RF motif selectively reduces the expression of ΔF508 CFTR in an EMC-dependent manner-potentially by promoting the premature termination of translation in response to misfolding.

**Figure 4.**
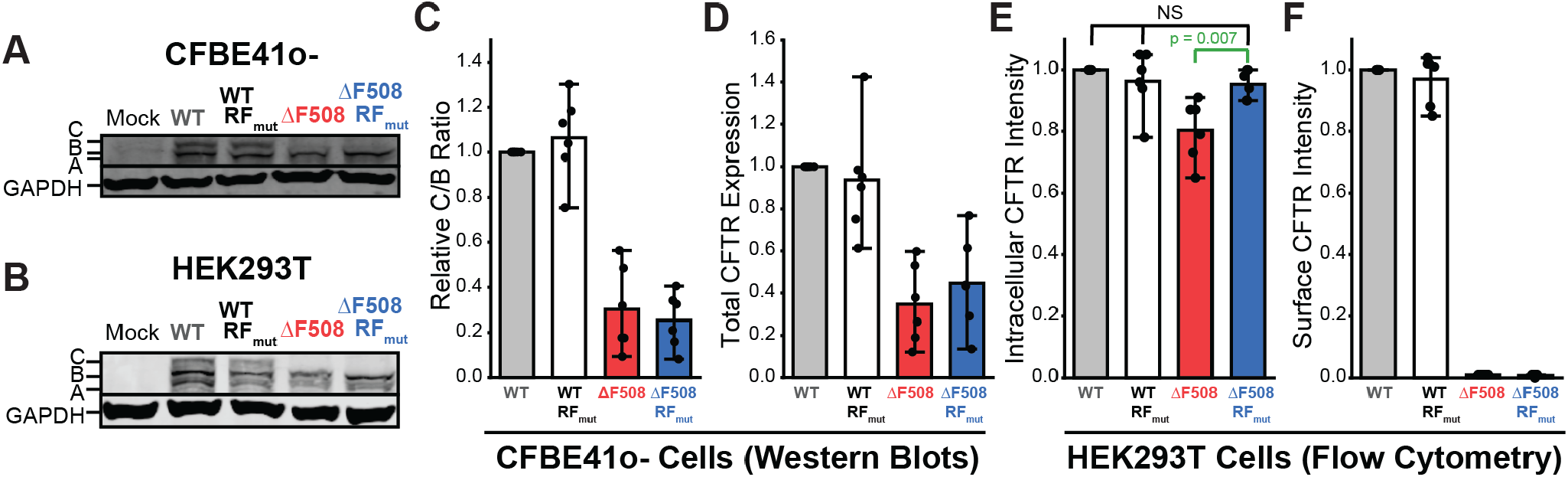
Impact of ribosomal frameshifting on CFTR expression. The expression and maturation of transiently expressed CFTR variants were compared in CFBE41o- and HEK293T cells by western blot and flow cytometry. A) A representative western blot depicting the relative abundance of the mature (band C) and immature (bands A & B) CFTR glycoforms of each indicated variant in CFBE41o-cells is shown. A GAPDH loading control is included for reference. B) A representative western blot depicting the relative abundance of the mature (band C) and immature (bands A & B) CFTR glycoforms of each indicated variant in HEK293T cells is shown. A GAPDH loading control is included for reference. C) A bar graph depicts the average C: B band intensity ratio relative to WT in CFBE41o-cells expressing the indicated variants as determined by western blot (n = 6). Error bars reflect the standard deviation. D) A bar graph depicts the average total CFTR intensity (C+B) relative to WT in CFBE41o-cells expressing the indicated variants as determined by western blot (n = 6). Error bars reflect the standard deviation. E) A bar graph depicts the average intracellular CFTR immunostaining intensity relative to WT among HEK293T cells expressing the indicated variants as determined by flow cytometry (n = 6). Error bars reflect the standard deviation. F) A bar graph depicts the average surface CFTR immunostaining intensity relative to WT among HEK293T cells expressing the indicated variants as determined by flow cytometry (n = 6). Error bars reflect the standard deviation.

### Impact of Ribosomal Frameshifting on CFTR Function

To determine how the RF motif impacts CFTR function, we compared the effects of the RF_mut_ modification on the ΔF508 and WT CFTR-mediated quenching of a cellular halide-sensitive yellow fluorescent protein (hYFP).^27^ Briefly, we generated a series of recombinant stable HEK293T cell lines that inducibly express a single CFTR variant off a bicistronic transcript containing a downstream internal ribosomal entry site (IRES) that produces the hYFP sensor and a fluorescent mKate reference fluorophore (Fig. 5A). We then used flow cytometry to track the CFTR-mediated quenching of hYFP fluorescence in relation to an mKate reference fluorophore at the single-cell upon addition of iodide (Fig. 5B). Cells expressing CFTR undergo an initial rapid hYFP quenching phase that occurs within the dead time for mixing that is followed by a slow exponential decay in the single-cell hYFP: mKate values (Fig. 5B). The observable hYFP quenching phase was 9-fold slower, on average, among cells expressing ΔF508 CFTR and was further delayed by a CFTR-specific inhibitor (CFTR(Inh)-172, Figs. 5B & S10), which suggests the observed quenching is rate-limited by CFTR channel conductance. Half-life values derived from a global fit of the slow hYFP quenching phase reveals that the RF_mut_ modification has minimal impact on WT CFTR but decreases the half-life of ΔF508 CFTR-mediated hYFP quenching by 46 ± 6% (Fig. 5B-C). These observations suggest the mutagenic disruption of the RF motif and its resulting impact on the CFTR interactome partially restores the function of ΔF508 CFTR. Interestingly, we find that the disruption of the RF motif does not result in an uptick in ΔF508 CFTR conductance in the context of EMC knockout cells (Fig. S11), which again suggests this feedback is EMC-mediated.

**Figure 5.**
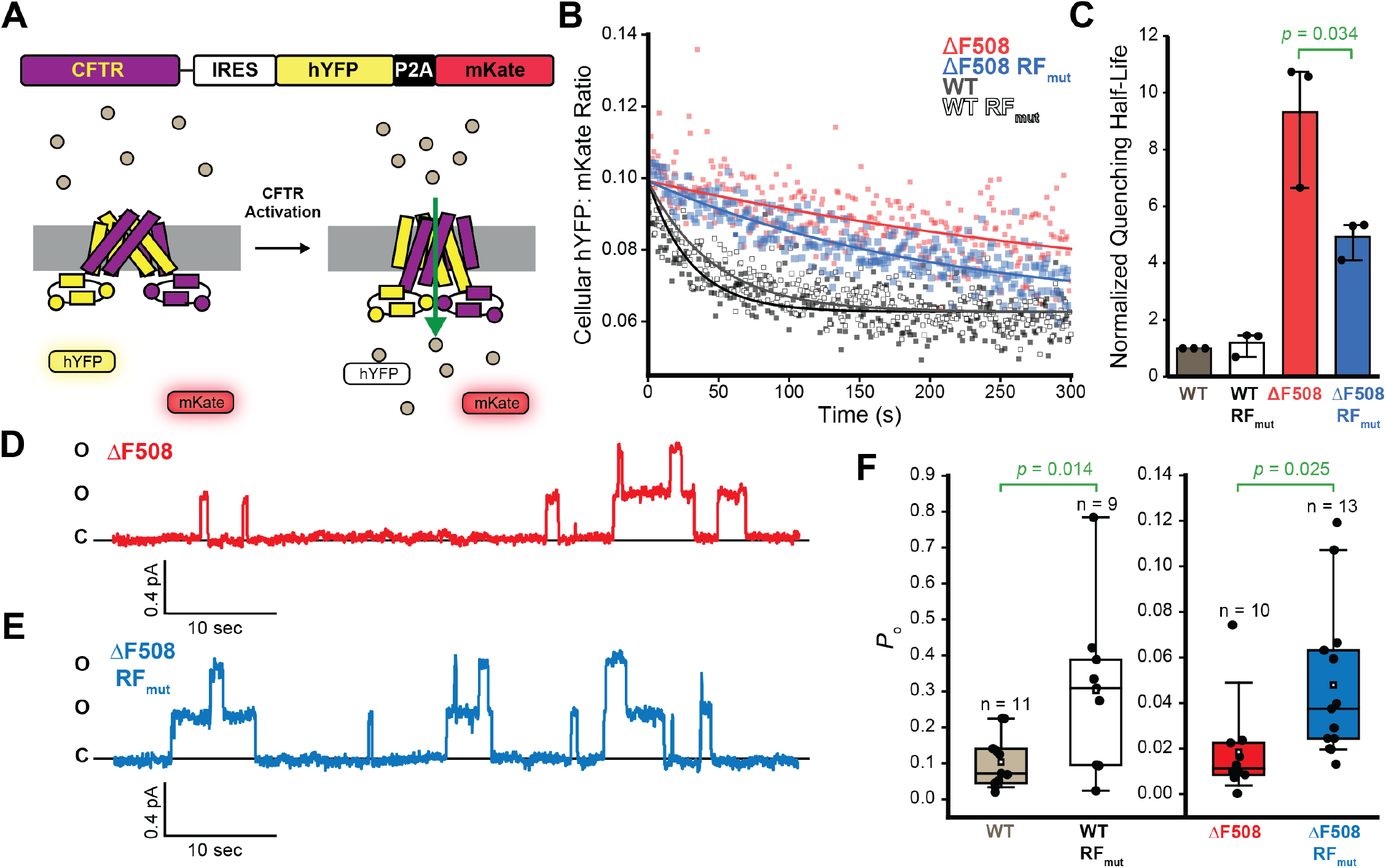
Impact of ribosomal frameshifting on CFTR function. The functional conductance of stably expressed CFTR variants was compared in HEK293T cells by measuring the time-dependent quenching of a halide-sensitive yellow fluorescent protein (hYFP). A) A cartoon depicts a diagram of the stably expressed genetic cassette (top) and a schematic for the CFTR activity assay (bottom). Upon activation, iodide ions flow through the CFTR channel and quench the hYFP fluorophore, which is monitored by the decrease of hYFP intensity relative to that of the fluorescent mKate standard. B) HEK293T cells stably expressing WT (white), WT RF_mut_ (gray), ΔF508 (red), and ΔF508 RF_mut_ (blue) were stimulated with 25 µM forskolin (0.06% DMSO Vehicle) to activate CFTR prior to measurement of the change in cellular hYFP: mKate intensity ratio measurements over time by flow cytometry. Cellular intensity ratios are plotted against the time and the global fits of the decay are shown for reference. C) A bar graph depicts the globally fit half-life for hYFP quenching for each variant. Values represent the average fitted values for each variant normalized relative to the WT value (n = 3). A *p*-value from a two-sample t-test is shown for reference. D) A representative current trace from an inside out macropatch of an HEK293T cell transiently expressing ΔF508 CFTR at 27° C is shown. E) A representative current trace from an inside out macropatch of an HEK293T cell transiently expressing ΔF508 CFTR RF_mut_ at 27° C is shown. F) A box and whisker plot depicts the statistical distributions of the fitted open probability (*P*_o_) values that were derived from a series of individual micropatch current traces that were collected from cells expressing WT (gray) or WT RF_mut_ (white) at 37°C and from cells expressing ΔF508 or ΔF508 RF_mut_ at 27°C. The upper and lower bounds of the boxes reflect the 75^th^ and 25^th^ percentile values and the upper and lower whiskers on both the boxes and the bars reflect the 90^th^ and 10^th^ percentile values, respectively. The midline and square reflect the median and average value, respectively. A Grubbs test was used to reject individual outliers and the *p*-values from two-sample student’s t-tests are shown for reference.

Disrupting the RF motif enhances hYFP quenching by ΔF508 CFTR without increasing its plasma membrane expression (Fig. 4F), which indicates this modification may improve the function of mature ΔF508 CFTR. We therefore compared the channel open probability (*P*_*o*_) of these variants in the context of inside-out macro- and/ or micropatches excised from the plasma membranes of transfected HEK293T cells. Briefly, we monitored single channel activity over time (8-13 min) in the presence of 1.5 mM ATP and Protein Kinase A (PKA, 124 U/mL) at a holding potential of 60 mV. We then estimated the number of channels in the patch and verified the identity of the channels with the sequential addition of a potentiator (VX-770) and a CFTR inhibitor (CFTR (inh)-172). Figures 5 D and 5E show representative current traces for patches containing ΔF508 and ΔF508 CFTR RF_mut_ CFTR, respectively. Currents across patches containing ΔF508 RF_mut_ CFTR generally feature more channel opening events relative to those containing ΔF508 CFTR (Fig. 5D-E). Overall, disruption of the RF motif increases the ΔF508 CFTR *P*_*o*_ by 1.6-fold (*p* = 0.025, Fig. 5F). The RF_mut_ modification has a similar impact on WT CFTR opening despite the fact that it has limited impact on the slow phase of cellular hYFP quenching (Fig. 5 C &F). Nevertheless, it is possible that the effect of the RF_mut_ modification of the single channel behavior of WT CFTR may be more pronounced within the initial rapid quenching phase that occurs within the mixing time. Regardless, these results confirm that the resulting modifications to the QC of nascent ΔF508 CFTR in the ER enhance the activity of the mature channel at the plasma membrane. Together, these cumulative findings reveal that the RF motif selectively modulates the assembly of ΔF508 CFTR in a manner that ultimately attenuates its expression and function.

### Impact of Ribosomal Frameshifting on the Pharmacological Rescue of ΔF508 CFTR

The functional expression of ΔF508 and various other CF variants can be partially restored by Trikafta, an FDA-approved cocktail of two correctors that stabilize the CFTR protein (VX-661 + VX-445) and a potentiator that activates it (VX-770).^28-30^ To determine whether ribosomal frameshifting potentially influences the effects of these compounds, we assessed whether the RF motif impacts the pharmacological rescue of ΔF508 CFTR. As expected, flow cytometry-based immunostaining measurements reveal that both WT and ΔF508 exhibit enhanced accumulation at the plasma membrane and within the secretory pathway upon stabilization by VX-661 + VX-445 (Fig. 6A-B). A western blot analysis also confirms that these correctors enhance the maturation of both variants (Fig. 6C, Fig S7). However, similar gains in expression and maturation were observed for WT-RF_mut_ and ΔF508-RF_mut_ (Fig. 6 A-C, Fig. S7), which demonstrates that the RF motif has minimal impact on ΔF508 expression upon stabilization by corrector molecules. Next, we assessed functional rescue of CFTR conductance using our hYFP quenching assay. For this purpose, the potentiator VX-770 was added in combination with the correctors VX-661 and VX-445 to evaluate the functional rescue by Trikafta. As a control, these quenching assays were also performed in the presence of a channel inhibitor (Fig. S8). Despite the comparable expression of ΔF508 and ΔF508-RF_mut_ in the presence of correctors, HEK293T cells expressing ΔF508-RF_mut_ that were treated with VX-661 + VX-445 + VX-770 exhibit a is 41 ± 5% decrease in hYFP quenching half-life relative to cells expressing ΔF508 CFTR under the same conditions (Fig. 6D). Together, these results suggest that, while stabilization by correctors minimizes the impact of the RF motif on ΔF508 CFTR expression, disrupting the RNA structure that promotes frameshifting ultimately enhances its functional rescue.

**Figure 6.**
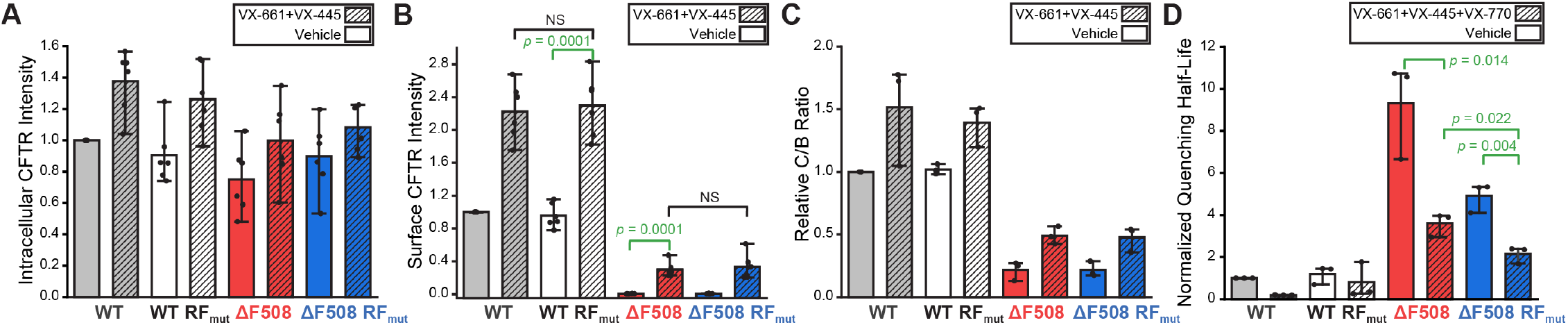
Impact of ribosomal frameshifting on the pharmacological rescue of ΔF508 CFTR. The effects of CFTR modulators on the expression and function of CFTR variants was determined in HEK293T cells. A) A bar graph depicts the average intracellular CFTR variant immunostaining intensity relative to WT among HEK293T cells treated for 16 hours with vehicle (open bars) or 3 μM VX-661 + 3 μM VX-445 (hashes) as determined by flow cytometry (n = 6). B) A bar graph depicts the average intracellular CFTR variant immunostaining intensity relative to WT among HEK293T cells treated for 16 hours with DMSO vehicle (open bars) or 3 μM VX-661 + 3 μM VX-445 (hashes) as determined by flow cytometry (n = 6). C) A bar graph depicts the average CFTR variant C: B band intensity ratios normalized relative to WT in HEK293T cells treated for 16 hours with vehicle (open bars) or 3 μM VX-661 + 3 μM VX-445 (hashes) as determined by western blot (n = 3). D) A bar graph depicts the average hYFP quenching half-life normalized relative to that of vehicle-treated WT for each variant as determined by flow cytometry (n = 3) in cells treated for 16 hours with vehicle (open bars) or 3 μM VX-661 + 3 μM VX-445 + 3 μM VX-770 (hashes) and pulsed with 25 µM forskolin. Measurements in the presence of vehicle from Figure 5C are shown for reference. Error bars reflect the standard deviation from three biological replicates in all cases. All treatments resulted in a final concentration of 0.06% DMSO (v/v).

## Discussion

The ΔF508 mutation promotes cotranslational CFTR misfolding and degradation in a manner that reduces its functional expression.^1^ Nevertheless, its proteostatic effects also stem from aberrant translational dynamics. It was previously shown that the impact of the ΔF508 mutation on the secondary structure of the CFTR transcript perturbs translation dynamics in a manner that compromises channel expression and function.^13,14^ Indeed, this mutation modifies interactions with over 20 translation factors and RNA processing proteins involved in CFTR biosynthesis.^5^ Moreover, the cotranslational misfolding of the nascent ΔF508 protein triggers interactions with the EMC in a manner that results in ribosome collisions and the premature termination of translation.^11^ However, it remains unclear how the cotranslational misfolding of the ΔF508 protein could modify the activity of the ribosome itself. Given that conformational transitions in the nascent chain can stimulate -1PRF,^15,16,31^ we searched the CFTR transcript for structural features that might promote ribosomal frameshifting. Our findings identified a slippery sequence (SSC) and stem-loop (SL2) that stimulate ribosomal frameshifting and the premature translational termination (Fig. 2). To determine how this RF motif impacts CFTR expression and function, we designed a series of silent mutations that maintain the native amino acid sequence in the 0-frame while disrupting these putative RNA structures (RF_mut_, Fig. 1). While this modification has minimal impact on WT CFTR assembly, disrupting the RF motif significantly remodels ΔF508 CFTR interactome (Fig. 3). These changes in ΔF508 assembly slightly increase the accumulation of ΔF508 CFTR within the secretory pathway but markedly enhance its functional conductance at the plasma membrane (Figs. 4 & 5). Finally, we show that disrupting the RF motif also enhances the pharmacological rescue of ΔF508 CFTR function (Fig. 6). Given that the disruption of this motif has minimal impact on the assembly, expression, or pharmacological profile of WT CFTR (Figs. 2-5), we conclude that this RNA structure stimulates RF and the premature termination of translation in response to cotranslational CFTR misfolding. Together, our findings suggest the RF motif acts as a “kill switch” that down-regulates the synthesis of defective proteins. Future investigations are needed to determine how this translational regulation influences the biosynthesis and pharmacological response of other known CF variants.

Interactome profiles suggest this RF motif may represent an assembly junction where the conformational state of the nascent chain modulates the interplay between translation and QC. We identify several ΔF508-specific chaperone interactions that are lost and others that are gained upon disruption of the RF motif (Table S1, Fig. 3). Moreover, weakening this RNA structure the context of the ΔF508 transcript significantly reduces the co-immunoprecipitation of several ribosomal proteins (Table S1, Fig. 3), which implies ribosomes typically stall within this region during ΔF508 translation. Based on these observations, we propose that the misassembly of nascent ΔF508 CFTR stimulates ribosomal frameshifting and directs the ribosome to a stop codon in the -1-frame (Fig. 7). Without the prompt intervention of ribosome quality control,^32^ this premature termination may cause trailing ribosomes to collide (Fig. 7). Such crosstalk may explain the link between cotranslational misfolding and ribosome collisions that occurs during pre-emptive QC.^11^ This interpretation is consistent with our observations that the effects of the frameshift motif are attenuated in EMC knockout cells (Figs. S9 & S11). Ribosomal pausing^11^ and/or heightened frameshifting during ΔF508 CFTR translation could potentially explain how this mutation enhances associations with UPF1 and other nonsense-mediated decay (NMD) proteins^5^ that typically promote the degradation of nonsense transcripts.^33,34^ Knocking down UPF1 and other associated RNA binding proteins partially restores ΔF508 expression,^5^ which until now has been a perplexing observation given that ΔF508 preserves the native reading frame. The stimulation of frameshifting by ΔF508 would promote premature termination via stop codons in the -1 frame. Such a link between ribosomal frameshifting and NMD has been previously described,^35^ and certain ribosome-associated molecular chaperones have been found to influence -1PRF.^36^ Nevertheless, to our knowledge, previous investigations have not established a direct role of ribosomal frameshifting in protein QC. We note that these interpretations remain speculative given that we are unable to compare ribosomal frameshifting efficiencies during CFTR biosynthesis and that our interactome measurements do not reveal a clear link to NMD. Additional investigations are needed to gain mechanistic insights into the interplay between cotranslational misfolding, ribosomal frameshifting, and RNA surveillance pathways.

**Figure 7.**
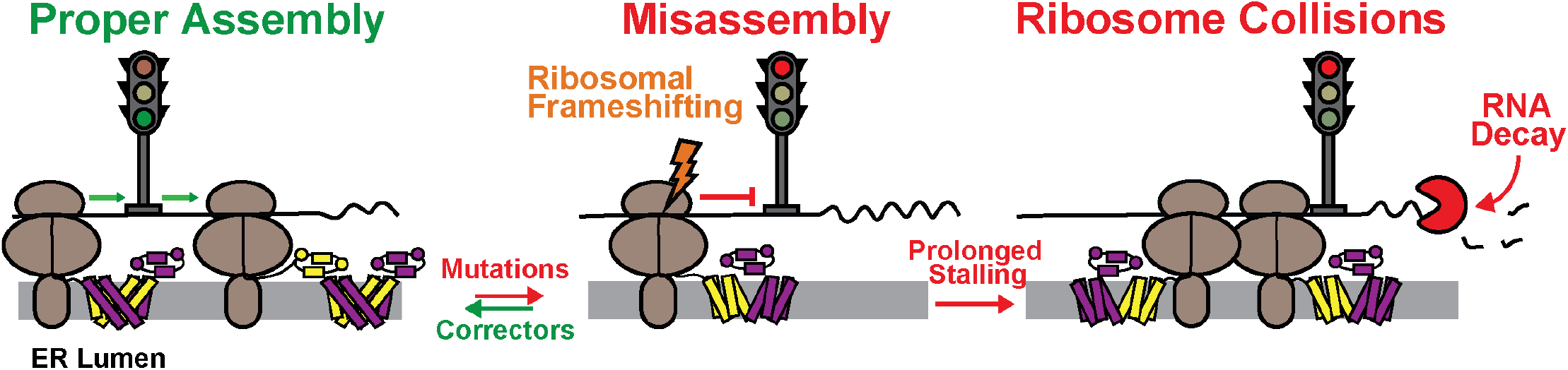
Proposed role of ribosomal frameshifting in the selective downregulation of ΔF508 CFTR biosynthesis. A cartoon depicts the proposed role of ribosomal frameshifting in CFTR proteostasis. Ribosomes pause at the ribosomal frameshift site at the point in which MSD1 and MSD2 undergo domain swapping. Synthesis proceeds with modest frameshifting when correct assembly occurs. However, failure to achieve the correct conformation and/ or chaperone interactions stimulates ribosomal frameshifting and the premature termination of termination of translation. Premature termination causes prolonged stalling that leads to collisions with trailing ribosomes and a failure to clear RNA surveillance proteins that trigger mRNA decay.

There are currently few examples of bona fide -1PRF sites within eukaryotic transcripts and the current evidence for such remains contentious.^24,37^ One key line of evidence that has cast doubt on the role of ribosomal frameshifting in eukaryotic gene regulation involves the general lack of the extended coding sequences within the alternative reading frames of eukaryotic genes- a classic indication of translational recoding in viral transcripts. Due to an abundance of stop codons in the -1 reading frame, ribosomal frameshifting in eukaryotes should most often result in premature termination of translation- an outcome that ribosomes have generally evolved to avoid. Nevertheless, premature termination and/ or the destabilization of the transcript may be beneficial under conditions in which the nascent polypeptides begin to misfold. Cells routinely undergo swings in adaptive proteostasis where the products of translation and the pool of molecular chaperones they rely on must be remodeled on short time scales. Based on our findings, we propose that this putative connection between cotranslational misfolding, ribosomal frameshifting, and RNA surveillance pathways provides a means for cells to remodel the transcriptome in response to misassembly in order to prevent the accumulation of misfolded proteins. Additional investigations are needed to determine whether similar QC checkpoints are present in the transcripts of other proteins that are predisposed to misfolding. Such investigations may offer novel mechanistic insights into the molecular basis of a wide array of diseases of aberrant cellular proteostasis.

## Materials and Methods

### Molecular Biology

All DNA constructs were cloned using the NEBuilder Hifi DNA Assembly System (NEB, cat # E2621X). All site directed mutagenesis reactions were performed using PrimeSTAR HS DNA Polymerase (Takara Bio, cat #R010B). Plasmids used for transient transfections in HEK293T cells were prepared and purified using the Zymopure Midiprep Kit (Zymo Research, cat #D4201). Mutations to create the CFTR ΔPRF constructs were chosen using RNA structure predictions performed by the ViennaRNA RNAfold Web Server.

### Cell Culture

HEK293T cells were grown in Dulbecco’s modified Eagle’s medium (Gibco, Grand Island, NY) containing 10% fetal bovine serum (Corning, Corning, NY) and a penicillin (100U/ml)/streptomycin (100µg/ml) antibiotic supplement (Gibco, Grand Island, NY) in a humidified incubator containing 5% CO2 at 37°C. HEK293T cells used for interactome measurements were grown under similar conditions but with 1% L-glutamine (200 mM) supplementation. Plasmid DNA constructs were transiently expressed in HEK293T cells using Lipofectamine 3000 (ThermoFisher Scientific, cat #L3000015). Cells were dosed with CFTR modulators a day after transfection when applicable. Two days post-transfection, cells were washed with 1X phosphate buffered saline (AthenaES, Baltimore, MD) and harvested with 1X 0.25% Trypsin-EDTA (Gibco, Grand Island, NY).

CFBE41o-cells were grown in minimal essential medium (Gibco, Grand Island, NY) containing 10% fetal bovine serum (Corning, Corning, NY) and a penicillin (100U/ml)/streptomycin (100µg/ml) (Gibco, Grand Island, NY) antibiotic supplement (Gibco, Grand Island, NY) in a humidified incubator containing 5% CO2 at 37°C on culture plates coated with PureCol purified bovine collagen (Advanced BioMatrix, Carlsbad, CA). CFBE41o-cells were transfected using Lipofectamine 3000 (ThermoFisher Scientific, cat #L3000015). Two days post-transfection, cells were washed with 1X hepes buffered saline (Gibco, Grand Island, NY) and harvested with 1X 0.25% Trypsin-EDTA (Gibco, Grand Island, NY).

CFTR function measurements were carried out in recombinant stable cells were made from genetically modified HEK293T cells grown in 10 cm dishes in complete media as was previously described.^38,39^ Briefly these HEK293T cells contain a Tet-Bxb1-BFP “landing pad” where recombination results in the integration of a plasmid into the gDNA of the cell.^38^ These cells were co-transfected with plasmid and a Bxb1 recombinase expression vector using Fugene 6 (Promega, Madison, WI). Doxycycline (2µg/mL) was added one day after transfection and the cells were grown for 3 days at 33°C. The cells were then incubated at 37°C for 24 hours prior to passaging for further experiments.

### Mass Spectrometry-Based Interactome Profiling

A pcDNA5 vector containing untagged CFTR with a CMV promoter was used in the interactome profiling experiments. Co-immunoprecipitation (co-IP) of CFTR bound with interactors was carried out as described previously.^25,40^ Briefly, cell lysates were normalized by dilution to a common protein concentration at 1 mL total volume and pre-cleared with 4B Sepharose beads (Sigma) at 4°C for 2 h while rocking. Precleared lysates were then immunoprecipitated with Protein G beads covalently crosslinked to 24-1 antibody (6 mg antibody/mL of beads) overnight at 4°C while rocking. Beads were washed three times with TNI buffer, twice with TN buffer, all buffer was removed with a needle and then frozen at –80°C for 1.5 hours. Proteins were then eluted off the beads while shaking at 750 RPM at 37°C for 30-60 min with elution buffer (0.2 M glycine, 0.5% IGEPAL CA-630, pH 2.3). The elution step was repeated once and combined to immediately neutralize with a 10:1 ratio of 1 M ammonium bicarbonate solution.

MS sample preparation of co-IP samples were performed as described previously.^25,26^ Briefly, samples were chloroform/methanol precipitated, rinsed with methanol, dried, and reconstituted in 1% Rapigest SF (Waters). Each sample was reduced with 5 mM TCEP (Sigma), alkylated with 10 mM iodoacetamide (Sigma), and digested with 0.5 μg of trypsin (Sequencing Grade, Promega, or Pierce) in 50mM HEPES (pH 8.0) at 37°C with shaking for a least 10 hours. Digested peptides diluted to 60 μL with water, labeled with 40 uL of TMT pro 16plex reagents (Thermo Fisher) for 1 hour, and quenched with 10% w/v ammonium bicarbonate. The sixteen TMT-labeled samples were pooled, acidified with MS-grade formic acid (Sigma), concentrated using a SpeedVac (Thermo Fisher), resuspended in 1500 μL of buffer A (95% water, 4.9% acetonitrile, and 0.1% formic acid), and loaded onto a triphasic MudPIT column. An Exploris 480 (Thermo Fisher) mass spectrometer equipped with an UltiMate3000 RSLCnano System (Thermo Fisher) was used for LC-MS/MS analysis as described previously.^25,26^

Proteome Discoverer 2.4 was used for peptide identification and TMT-based quantification as described previously.^26^ MS/MS spectra were searched using SEQUEST against a UniProt human proteome database (released 03/25/2014) using a decoy database of reversed peptide sequences. The following parameters were employed: 0.02 Da fragment mass tolerance, 10 ppm peptide precursor tolerance, six amino acid minimum peptide length, trypsin cleavage with a max of two missed cleavages, static cysteine modification of 57.0215 Da (carbamidomethylation), and static N-terminal and lysine modifications of 304.2071 Da (TMT pro 16plex). Percolator was used to filter SEQUEST search to minimize the peptide false discovery rate to 1% and require a minimum of two peptides per protein identification. Reporter Ion Quantification processing node in Proteome Discoverer 2.4 was used to quantify TMT reporter ion intensities and a summation was performed for peptides in the same protein.

### Interactor filtering and data analysis

A total of 5-6 sets of samples per condition (WT, WT-RF_mut_, ΔF508, and ΔF508-RF_mut_) were analyzed via LC-MS/MS over 3 separate mass spectrometry runs. Co-immunoprecipitated interactors were compared against GFP transfection control to obtain log2 fold enrichment and adjusted p-value. To filter for statistically significant interactors of CFTR, log2 fold change over 1 and adjusted p-value below 0.05 were used as cutoffs. Interactors in each condition were filtered and combined to generate a master list. This list was queried against the Contaminant Repository for Affinity Purification (CRAPome 2.0)^41^ to remove proteins with a nonspecific detection threshold of over 500 reported experiments. The grouped abundance values of these interactors in each condition were log2 transformed to yield the consensus log2 grouped abundance for each protein. These values were normalized to WT CFTR log2 grouped abundance to standardize protein abundances across conditions.

### Hierarchical clustering and network analysis

Hierarchical clustering of proteins in the interactor master list was performed with a custom R script as described previously.^26^ Briefly, the log2 grouped abundance that were normalized against WT CFTR were converted to an Euclidean distance matrix and clustered using Ward’s minimum variance method. Network plotting of each cluster was performed using the String database (string-db.org) with interaction score set at medium confidence (0.400). Interactors not found in the String database were added manually to the network plots as isolated nodes. K-means clustering was performed for each hierarchical cluster.

### Luciferase-Based Ribosomal Frameshifting Measurements

Ribosomal frameshifting measurements were made using a previously described dual luciferase reporter system that was modified to avoid aberrant splicing artifacts.^24^ Briefly, all dual luciferase reporters were generated from a pJD2044 plasmid containing firefly and renilla luciferase and a CMV promoter. Potential cryptic splice sites in the renilla luciferase gene were identified using the human splicing finder webtool and were removed using a series of synonymous codon mutations (246T>C, 249G>A, 888T>G, and 891G>A).^42^ A 138 bp fragment of the CFTR transcript (3963-4101) was inserted between the 5′ renilla luciferase (0-frame) and 3′ firefly luciferase (−1-frame) genes. P2A linkers were inserted both upstream and downstream of the CFTR insert. HEK293T cells were plated in six well dishes one day prior to transfection of reporter constructs using Lipofectamine 3000 (ThermoFisher Scientific, cat #L3000015). Two days post-transfection, cells expressing dual luciferase reporter constructs were harvested and lysed using the Passive Lysis Buffer provided in the Dual-Luciferase Reporter Assay System (Promega, cat #E1980). Relative activities of the firefly and renilla luciferase domains in the clarified lysate were then measured using the Dual-Luciferase Reporter Assay System on a Synergy Neo2 plate reader (Biotek, Winooski, VT) in accordance with the manufacturer’s instructions. To determine the –1 ribosomal frameshifting efficiency the firefly: renilla luciferase ratio was normalized relative to that of a construct lacking the insert as was previously described.^43^

### Western Blot-Based CFTR Expression Measurements

Western blotting was used to evaluate the relative abundance and maturation of CFTR glycoforms in both HEK293T and CFBE41o-cells. Cells were grown in 6 cm dishes and transfected with Lipofectamine 3000 (ThermoFisher Scientific, cat #L3000015). Un-tagged CFTR variants were transiently expressed from a pcDNA5 vector featuring a CMV promoter. Cells were passively lysed two days post-transfection by rocking the cells at 4° C in a lysis buffer containing 25 mM Tris (pH 7.6), 150 mM NaCl, 1% Triton X-100, and 2mg/ mL of a protease inhibitor cocktail (Roche Diagnostics, Indianapolis, IN). Total protein concentrations within the cleared lysates were then determined using a detergent-compatible Bradford assay (Pierce Biotechnology, Waltham, MA). Lysates were diluted into SDS-PAGE sample buffer and heated to 37°C for 30 min prior to loading 65µg of total protein from each sample onto a 7.5% SDS-PAGE gel. Proteins were then separated by electrophoresis and transferred onto a PVDF membrane overnight at 4°C. Following transfer, the membrane was blocked in a 5% milk solution in TBST for 2 hours at room temperature. After two hours the lower part of the membrane was cut to isolate the GAPDH loading control bands. The membranes were incubated in a wash solution containing either a mouse anti-CFTR antibody (1:1,000 dilution AB217, CFTR Antibody Distribution Program, Chapel Hill, NC) or a mouse anti-GAPDH antibody (1:10,000 dilution ab8245, Abcam, Cambridge, UK) overnight at 4°C. Both CFTR and GAPDH containing membranes were then washed three times in TBST prior to incubation in a wash solution containing an IRDye 680RD-labeled goat anti-mouse secondary antibody (1:10,000 dilution, LI-COR Biosciences, Lincoln, NE) for two hours. Following incubation, the membranes were washed three more times in a TBST wash solution and allowed to dry for 30 min prior to imaging using an Odyssey CLx system (LI-COR Biosciences, Lincoln, NE). Band intensities were analyzed using either Image Studio V. 5.2 software (LI-COR Biosciences, Lincoln, NE) or Image J.

### Flow Cytometry-Based CFTR Expression Measurements

Flow cytometry was used to compare the plasma membrane and intracellular expression levels of CFTR mutants using a previously described approach.^44^ Briefly, CFTR variants were transiently expressed in HEK293T cells using a pcDNA5 vector featuring a CMV promoter, CFTR variants that were generated in the context of a previously described triple hemagglutinin (HA) tag in the fourth extra-cellular loop,^45^ and an internal ribosome entry site (IRES)-eGFP cassette. Cells were transfected in 6 cm dishes for 24 hours prior to treatment with either DMSO (vehicle) or 3µM VX-445 and VX-661. Cells were harvested 48 hours post transfection prior to immunostaining the plasma membrane CFTR for 30 min. with a DyLight 550-conjugated anti-HA antibody (ThermoFisher, Waltham, MA). Cells were then fixed, washed, and permeabilized using the Fix and Perm kit (Invitrogen, Carlsbad, CA). Intracellular CFTR was the immunostained for 30 minutes in the dark using an Alexa Fluor 647-conjugated anti-HA antibody (Invitrogen, Carlsbad, CA). Cells were washed and filtered prior to analysis of cellular fluorescence profiles using a BD LSRII flow cytometer (BD Biosciences, Franklin Lakes, NJ). Forward and side scatter measurements were used to gate for intact single cells. eGFP intensities (488 nm laser, 530/30 nm emission filter) were used to gate for positively transfected cells. Dylight 550 (561 nm laser, 582/15 nm emission filter) and Alexa Fluor 647 (640 nm laser, 670/30 nm emission filter) intensities were then measured for at least 10,000 positively transfected cells within each biological replicate. Data was analyzed using FlowJo software (Treestar, Ashland, OR).

### CFTR-Mediated hYFP Quenching Measurements

A fluorescent quenching functional assay based on a previously described method^27,46^ was used to compare the relative functional conductance of CFTR mutants. A pcDNA5 vector bearing an attB recombination site in place of its CMV promoter, untagged CFTR cDNA encoding each variant, and an IRES hYFP-P2A-mKate sensor cassette was used to generate a series of stable HEK293T cell lines that inducibly express each CFTR construct as previously described.^38,39^ Recombinant HEK293T cells stably expressing CFTR were grown in six well dishes and dosed with either DMSO (vehicle) or 3µM VX-445 and VX-661 48 hours after plating. Following 18 hrs. of treatment with correctors, the cells were harvested with trypsin, washed once in PBS containing 5 mM EDTA (pH 7.4), washed twice in PBS containing 137 mM sodium gluconate (pH 7.4), then resuspended in PBS containing 137 mM sodium gluconate (pH 7.4), 25 µM Forskolin, 50 µM Genestein, and 10 µM VX-770. Cells were incubated in the resuspension media for 10 minutes prior to a final wash and resuspension in PBS containing 137 mM sodium gluconate (pH 7.4) prior to analysis. hYFP quenching was initiated with the addition of 25 mM sodium iodide immediately prior to the analysis of single-cell forward light scattering, side light scattering, hYFP fluorescence (488 nm laser, 530/30 nm emission filter), and mKate fluorescence (561 nm laser, 610/20nm emission filter) intensity values as a function of time using a BD LSRII flow cytometer (BD Biosciences, Franklin Lakes, NJ). Data was analyzed using FlowJo software (Treestar, Ashland, OR). OriginPro 2019 (Origin, Northampton, MA) was used to globally fit the decay of single-cell hYFP: mKate ratios as a function of time using the following exponential decay function:

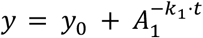

where *y* is the hYFP: mKate fluorescence intensity ratio at time *t, y*_*0*_ is the baseline hYFP: mKate fluorescence intensity ratio at *t* → ∞, *A*_*1*_ is the amplitude of the signal, and *k*_*1*_ is the observed rate constant for the exponential decay. The value of *y*_*0*_ was fixed in the global fit to an experimentally determined value for the hYFP: mKate intensity ratio of cells prior to their treatment with sodium iodide. The quenching half-life was calculated from the globally fit parameters using the following equation:

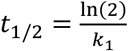

where *t*_*1/2*_ is the half-life and *k*_*1*_ is the globally fit rate constant.

### Patch Clamp Measurements

Patch clamp recording was used to estimate and compare the channel open probability (P_o_) of ΔF508 CFTR and ΔF508-RF_mut_ CFTR. Briefly, unitary currents were recorded in excised, inside-out patches from HEK293T cells transiently transfected with either construct. Both ΔF508 and ΔF508-RF_mut_ expressing cells were incubated at 27° C for 24-48 hours before patch experiments. Patch pipettes were pulled from Warner Instrument G85150T-3 glass to a tip resistance of 4-12 MΩ. Both the pipette and bath solutions were symmetrical and contained 140 mM N-Methyl-D glucamine, 3 mM MgCl2, 1 mM EGTA and 10 mM TES (pH 7.3). CFTR channels were activated by 1.5 mM Mg-ATP and 123 U/ml of recombinant Protein Kinase A (PKA, Promega, Madison, WI) in the cytosolic bath. Patches were held at +60 mV for unitary current recordings which were analog filtered at 200 Hz and then digitally filtered at 10 Hz with Clampfit 9.2 software (Axon Instruments, San Jose, CA). Data acquisition and analysis were performed using pCLAMP 9.2 software. All patch-clamp experiments were performed at room temperature.

*P*_o_ values of ΔF508 CFTR and ΔF508-RF_mut_ CFTR were determined from patches as described previously.^47,48^ Recordings containing fewer than 8 simultaneous openings prior to the addition of VX-770 (200 nM) were selected for analysis. The number of channels in each patch (*N*) and the single channel *P*_o_ for that patch (*NP*_o_) were estimated using Clampfit 9.2. The *P*o value of channels under control conditions was then estimated by dividing the total number of channels (*N*) in that patch which was estimated from the steady-state macroscopic current measured after potentiator addition; i.e., as *N* = *I*/*i* where *I* is the steady-state macroscopic current measured after potentiator addition and *i* is the unitary current at the holding potential of 60 mV (0.4 pA). This provides a minimal estimate of *N* and a maximal estimate of *P*_o_ before potentiator addition due to the underlying assumption that the *P*_o_ for potentiator-activated channels approaches unity.

### Generation and Validation of CRIPSR EMC6 Knockout Cells

Cas9 nuclease, a 5’ ATTO-labelled trans-activating CRISPR RNA (tracrRNA), and a Cas9 nuclease CRISPR guide RNA (gRNA) targeting the *EMC6* gene, the disruption of which was previously shown to destabilize the EMC complex,^49^ were purchased through IDT using the Alt-R CRISPR Cas9 system (IDT, Newark, NJ). Prior to introduction into HEK293T cells, the gRNA:tracrRNA duplex was prepared by mixing a 1:100 dilution of the gRNA and tracrRNA, and incubating the solution to 95°C for 5 minutes. The solution was allowed to cool to room temperature. The Cas9 nuclease was also diluted 1:62 with PBS (pH 7.2). A reverse transfection was performed using the Lipofectamine CRISPRMAX transfection reagent (ThermoFisher, Waltham, MA) to genomically modify the cells. Briefly, the ribonucleoprotein complex containing the gRNA:tracrRNA duplex and Cas9 nuclease was incubated at room temperature for 5 minutes. Cells were added to the CRISPRMAX transfection reagent containing the generated RNP complex and nuclease in a 96-well dish. Cells were incubated at 37°C for 48 hours prior to single-cell sorting on a BD FACSAria-II cell sorter (BD Biosciences, Franklin Lake, NJ) based on ATTO 550 fluorescence. Single cells were recovered and grown to confluency in complete Dulbecco’s modified Eagle medium (Gibco, Carlsbad, CA) supplemented with 10% fetal bovine serum (Corning, Corning, NY) and penicillin (100 U/ml)/ streptomycin (100 μg/ml) (complete media) before subsequent validation and confirmation of genome editing.

CRISPR genome editing of clones were validated with a T7E1 endonuclease assay, western blotting, and Sanger sequencing. Briefly for the T7E1 assay, gDNA from the sorted, grown cells were extracted using the GenElute Mammalian Genomic DNA Miniprep kit (Sigma-Aldrich, St. Louis, MO). A PCR of the EMC6 region was performed with the gDNA as the template and the PCR product was purified before hybridization with the T7E1 endonuclease (New England Biolabs, Ipswitch, MA). This amplified region was also sent out for Sanger sequencing validation. Post-T7E1 hybridization, a PCR cleanup of the reaction was run on a 2% agarose gel to observe banding patterns of genome edited cells. Western blots were run with a protocol previously described with a few changes described henceforth.^50^ Confluent cells were lysed via sonication using a RIPA lysis buffer containing 150 mM NaCl, 1% Nonidet P-40, 0.5% DOC, 0.1% SDS, and 2 mg/ mL of a protease inhibitor cocktail (Roche Diagnostics, Indianapolis, IN) in 50 mM Tris (pH 7.4). Total protein concentrations were determined using a detergent-compatible Bradford assay (Pierce Biotechnology, Waltham, MA). Lysates were then diluted into SDS-PAGE sample buffer and 20 μg of total protein from each sample was loaded onto a 12% SDS-PAGE gel. Proteins were then separated by electrophoresis and transferred onto a PVDF membrane. Membranes were blocked using the Intercept (TBS) Blocking Buffer (LI-COR Biosciences, Lincoln, NE) prior to incubating the membrane in a solution containing a rabbit anti-EMC6 antibody (1:300 dilution anti-EMC6 with 0.1% Tween 20, Abcam, Cambridge, UK). Membranes were washed three times with TBST buffer prior to incubating the membrane in a wash solution containing an IRDye 680RD-labeled goat anti-rabbit secondary antibody (1:5,000 dilution, LI-COR Biosciences, Lincoln, NE). The membrane was then washed three more times in TBST prior to fluorescent imaging using an Odyssey CLx system (LI-COR Biosciences, Lincoln, NE). Loading controls were collected for each blot by stripping the membrane of the primary and secondary antibody probes and repeating the protocol using the same protocol, except with a anti-Cyclophilin B primary antibody (1:1,500 dilution, Sigma, St. Louis, MO) was used in place of the anti-EMC6 antibody.

## Acknowledgements

We thank Suchetana Mukhopadhyay, Jonathan Dinman, and Jeff Brodsky for helpful critiques of our initial findings. We thank Christiane Hassel for technical input and assistance. We also acknowledge the support of the Indiana University Flow Cytometry Core Facility and the Indiana University Center for Genomics and Bioinformatics. This research was supported in part by grant from the National Institute of General Medical Sciences to J. P. S. (R01GM138845).

