## Supplemental Materials for "Ribosomal Frameshifting Selectively Modulates the Assembly, Function, and Pharmacological Rescue of a Misfolded CFTR Variant"

**Contents:**

-Figure S1
-Figure S2
-Figure S3
-Figure S4
-Figure S5
-Figure S6
-Figure S7
-Figure S8
-Figure S9
-Figure S10
-Figure S11
-Table S1

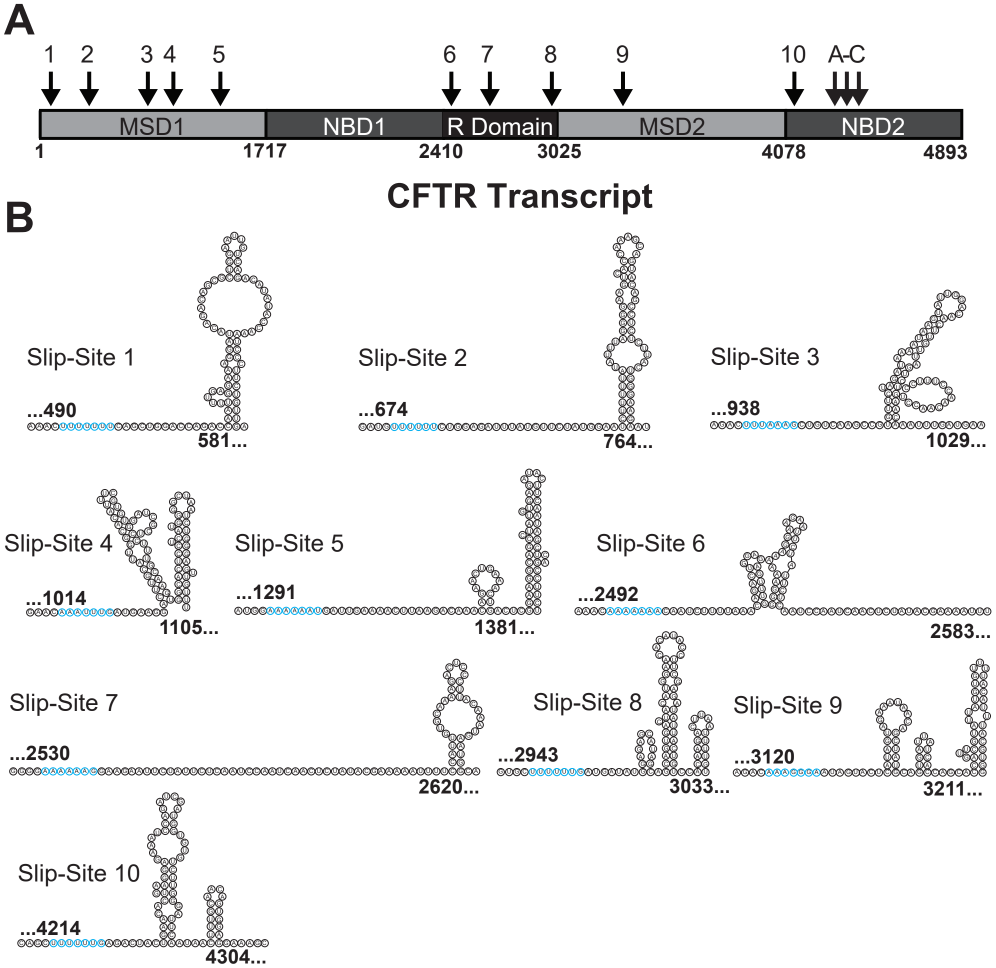


**Figure S1. Potential Slip Sites in CFTR.** The relative positions of various canonical and near-canonical $X_1$ $XXY_4$ $YYZ_7$ sites within the CFTR trasncript along with downstream secondary structure predictions are shown. (A) A schematic depicts the nucleotide position of the protein domain boundaries and potential slip-sites within the CFTR transcript. (B) Efficient frameshifting typically requires the formation of stable RNA secondary structure 5-8 basepairs downstream of the slippery sequence. We therefore utilized the Vienna RNAfold program to predict the mRNA secondary structure within the 75 nucleotides that lie 5 basepairs downstream of each putative slippery site. Cartoons depict the most stable predicted secondary structure at each site in relation to the slippery sequence (blue) for each potential slip-site not characterized in this work.

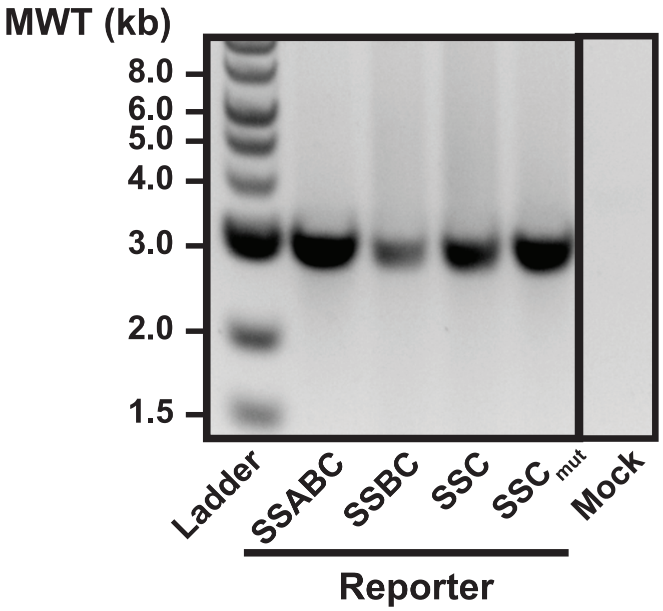


**Figure S2. Reverse Transcriptase PCR of CFTR Dual Luciferase Reporters.** CFTR frameshift remporters were transiently expressed in HEK293T cells prior to the extraction of the total cellular mRNA and the amplification of each reporter cDNA with reverse transcriptase. The products of these reactions are shown on an image of a representative 1% agarose gel. Reactions produced a clean, strong band in the expected molecular weight range (~3 kb) in all cases, which indicates the reporter constructs generate the expected transcript without any significant alternative splicing.

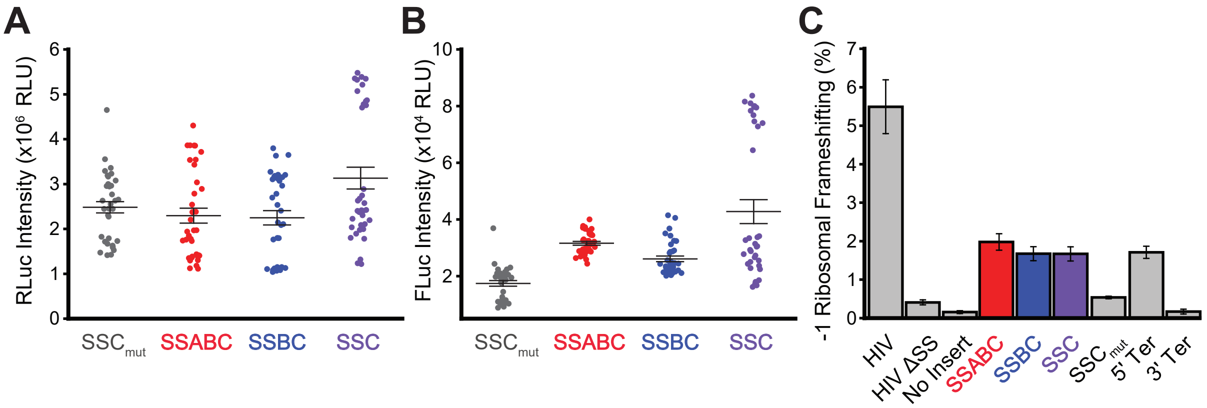


**Figure S3. Ribosomal frameshifting reporter measurements in HEK293T Cells.** The efficiency of ribosomal frameshifting during the translation of a structured region within the CFTR transcript was assessed in HEK293T cells using a bicistronic dual luciferase reporter system. (A, B) Dot plots indicate the raw RLuc and FLuc luminescence intensities within the lysates of HEK293T cells transiently expressing these constructs, respectively. Results were taken from three biological replicates with 4 transfections per sample each. Central hashes represent the average intensities and whiskers reflect their standard deviation. (C) A bar graph depicts the average -1 ribosomal frameshifting efficiencies for each construct in HEK293T cells. Error bars reflect the standard deviation to serve as a measure of precision.

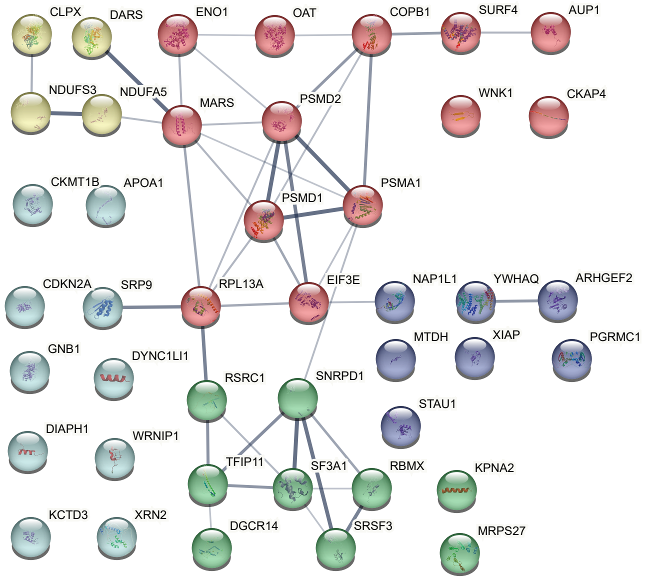


**Figure S4. Network model of the CFTR interactors within heirachical cluster 2.** A network map depicts the relationships between interactors in cluster 2. Lines indicate known protein-protein interactions in the String database (human). Line widths indicate the strength of data support. The colors of nodes reflect the identity of sub-clusters from K-means clustering of the interactors within hierarchical cluster 2.

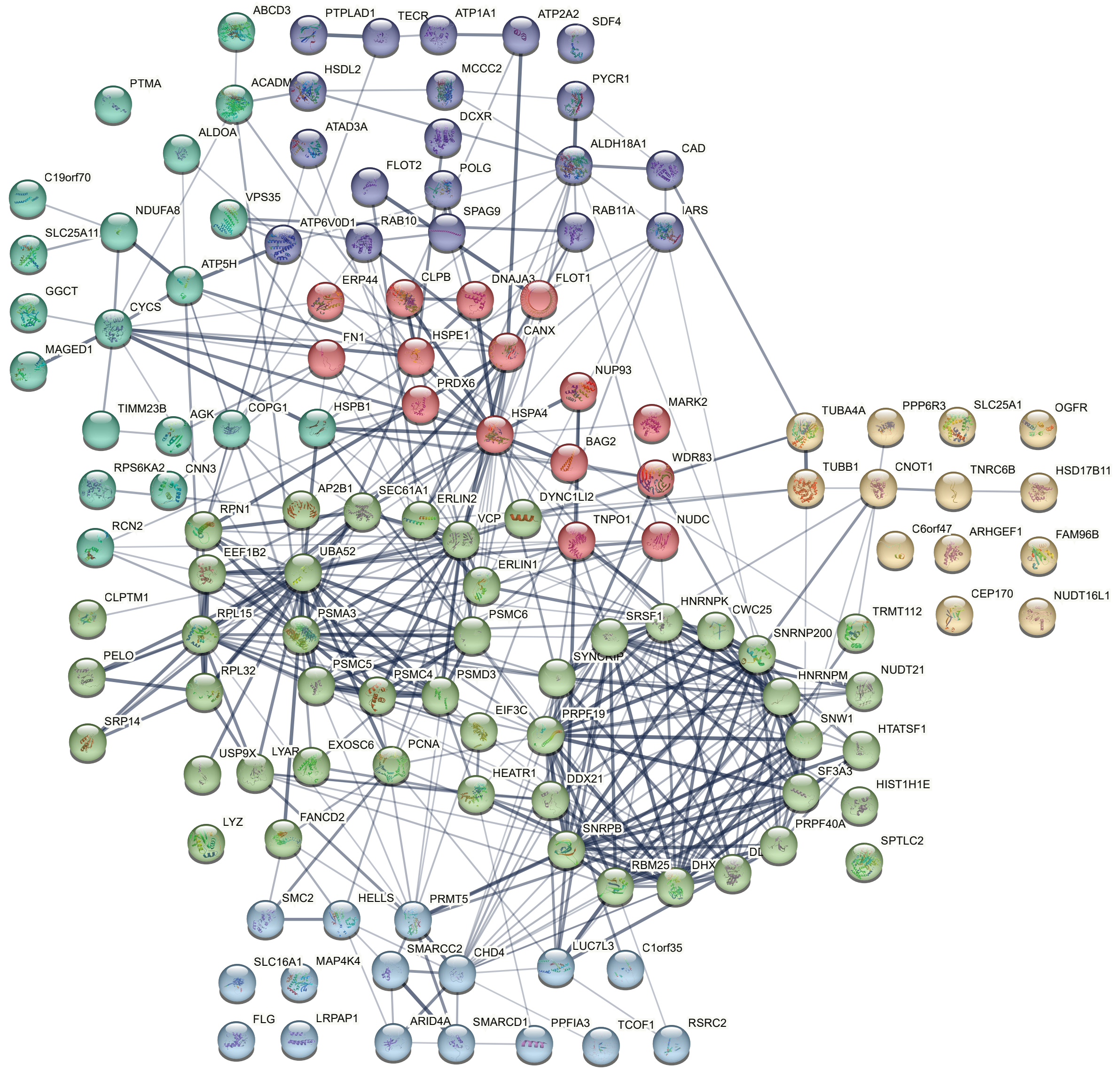


**Figure S5. Network model of the CFTR interactors within heirachical cluster 3.** A network map depicts the relationships between interactors in cluster 3. Lines indicate known protein-protein interactions in the String database (human). Line widths indicate the strength of data support. The colors of nodes reflect the identity of sub-clusters from K-means clustering of the interactors within hierarchical cluster 3.

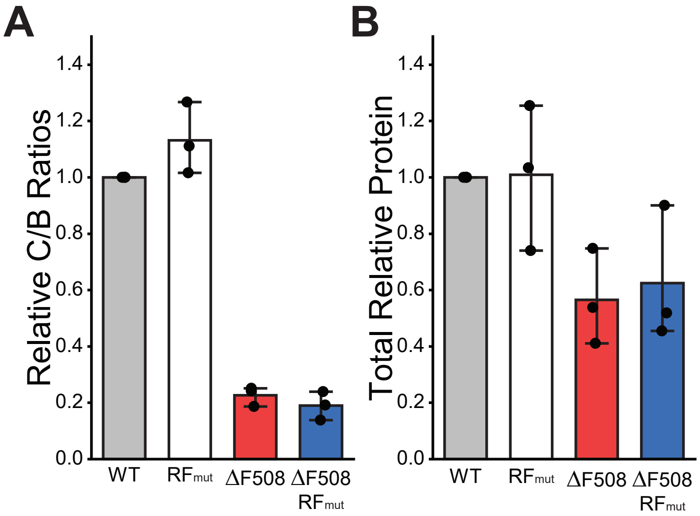


**Figure S6. Impact of ribosomal frameshifting on CFTR expression in HEK293T cells.** The expression and maturation of transiently expressed CFTR variants are quantified in HEK293T cells by western blot. A) A bar graph depicts the average C: B band intensity ratio relative to WT in HEK293T cells expressing the indicated variants as determined by western blot (n = 3). Error bars reflect the standard deviation. B) A bar graph depicts the average total CFTR intensity (C+B) relative to WT in HEK293T cells expressing the indicated variants as determined by western blot (n = 3). Error bars reflect the standard deviation.

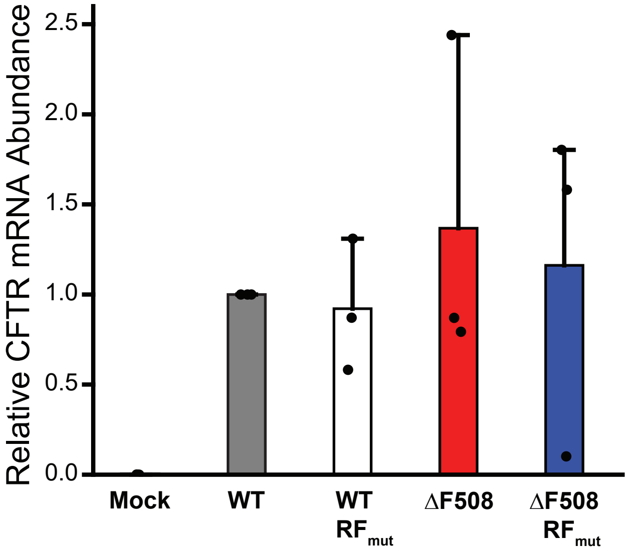


**Figure S7. Relative abundance of CFTR transcripts in transiently transfected CFBE 41 o- cells.** Quantitative PCR (qPCR) was used to compare the abundance of the CFTR transcript in CFBE41o- cells transiently expressing WT (gray), WT $RF_{mut}$ (white), ΔF508 (red), and ΔF508 $RF_{mut}$ (blue).Cells were harvested two days after transfection prior to the extraction of cellular RNA. Cellular mRNA were then reverse-transcribed into DNA and real-time PCR was used to track TaqMan probe amplification from the cDNA for CFTR and an endogenous control transcript (ACTB). ΔCt values for each variant were used to quantify the abundance of each transcript relative to that of WT (RQ), which are plotted in the bar graph above. Points represent individual values from three biological replicates. Bars report the average values and the whiskers show the 90th percentile value. A mock sample that was transfected without the expression vector is shown for reference. The results reveal that transient transfections generate a high degree of variability in transfection efficiency, which is likely to obscure any deviations in transcript stability.

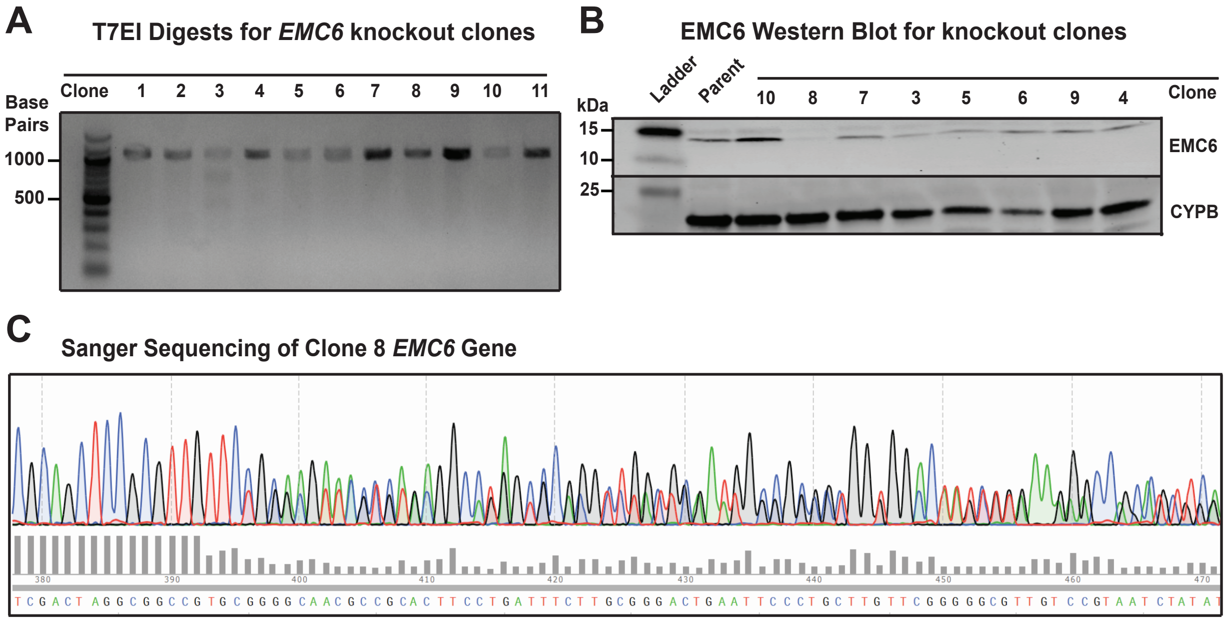


**Figure S8. HEK293T EMC6 Knockout Cell Line Validation.** Select HEK293T clones that were isolated following a CRISPR-mediated *EMC6* knockout were characterized using a T7E1 endonuclease assay, western blotting, and Sanger sequencing of the *EMC6* gene. A) Following Cas9-mediated gene editing, the *EMC6* gene was amplified from the genomic DNA of each clone then melted and re-annealed prior to the digestion with T7E1 endonuclease. Digestion of the band is indicative of mismatches arising from heterogeneous DNA edits/ repairs. B) A western blot for the endogenous EMC6 protein is shown for several of the isolated clones. An unedited HEK293T cell line is shown for reference and a Cyclophilin B (CYPB) loading control is included. C) A Sanger sequencing chromatogram is shown for the *EMC6* gene from Clone 8. Quality scores for individual DNA bases, which reflect the heterogeneity of the underlying sequence, are shown for reference. Sequencing data downstream of base 392 suggests one or more fameshift edits occured in this clone, which was used for the experiments detailed herein.

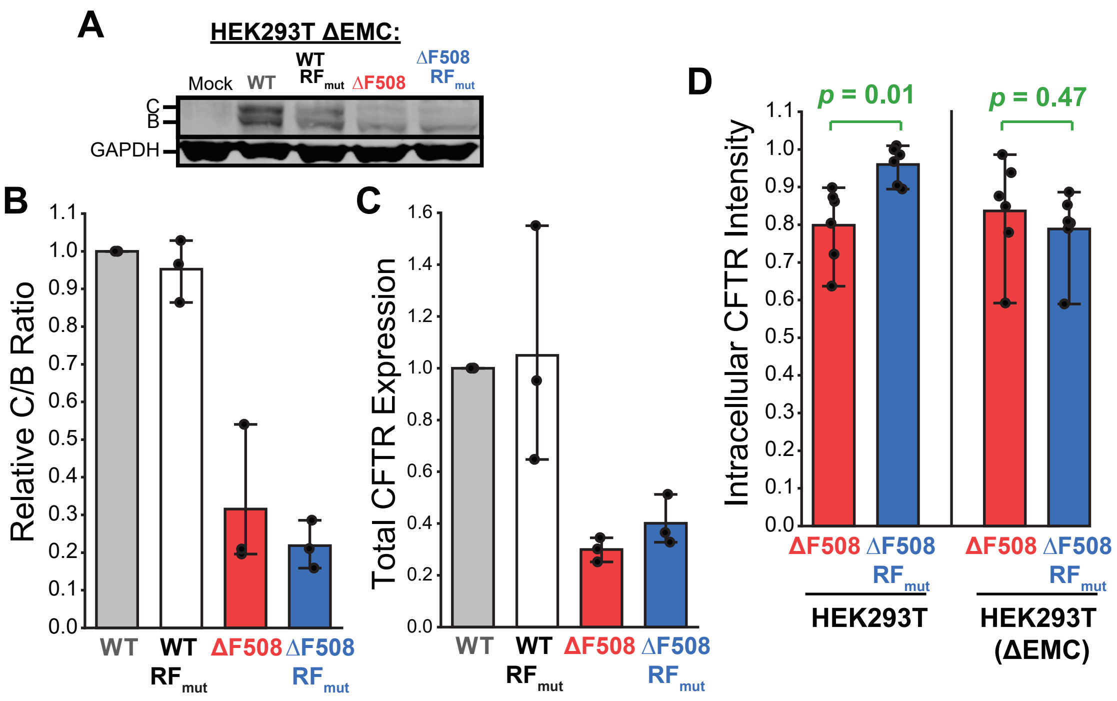


**Figure S9. CFTR Expression in *EMC6* Knockout Cells.** The expression and maturation of transiently expressed CFTR variants were assessed in the context of an *EMC6* knockout HEK293T cell line by western blot and flow cytometry. A) A representative western blot depicting the relative abundance of the mature (band C) and immature (band B) CFTR glycoforms of each indicated variant in is shown. A GAPDH loading control is included for reference. B) A bar graph depicts the average C: B band intensity ratio relative to WT in expressing the indicated variants as determined by western blot (n = 3). Error bars reflect the standard deviation. C) A bar graph depicts the average total CFTR intensity (C+B) relative to WT for the indicated variants as determined by western blot (n = 3). Error bars reflect the standard deviation. D) A bar graph depicts the average intracellular CFTR immunostaining intensity relative to WT among HEK293T cells (left) and *EMC6* knockout cells (right) expressing the indicated variants as determined by flow cytometry (n = 6). Error bars reflect the standard deviation. *p*-values from a two-sample t-test are shown for reference.

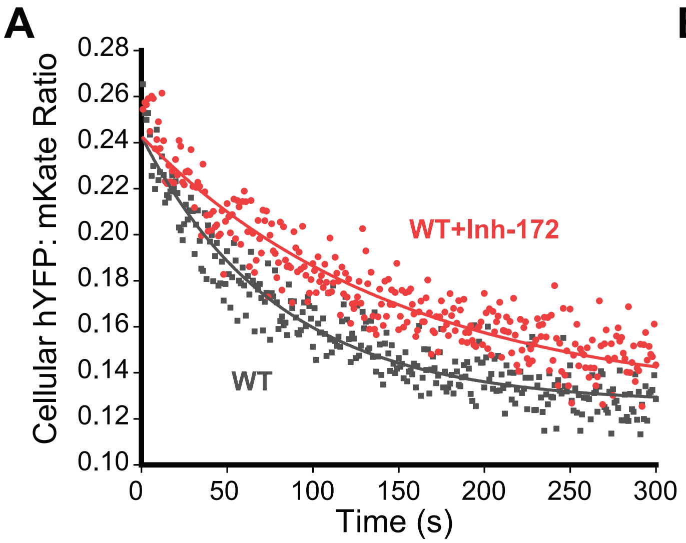


**Figure S10. Impact of CFTR Inh-172 on the Observable hYFP Quenching Kinetics.** The imact of the CFTR-specific inhibitor 172 on the kinetics of halide-sensitive yellow fluorescent protein (hYFP) quenching upon activation of stably expressed CFTRs in HEK293T cells was characterized by flow cytometry. A) HEK293T cells stably expressing WT CFTR were stimulated with 25 μM forskolin (0.06% DMSO Vehicle) to activate CFTR prior to measurement of the change in cellular hYFP: mKate intensity ratio measurements over time in the presence (red) and absence (gray) of 10 μM Inh-172 by flow cytometry. Single cell intensity ratios are plotted against the time and the fits for a single-exponential decay function are shown for reference. The fitted quenching half-life value increases 89% in the presence of inhibitor. A) HEK293T cells stably expressing ΔF508 CFTR were stimulated with 25 μM forskolin (0.06% DMSO Vehicle) to activate CFTR prior to measurement of the change in cellular hYFP: mKate intensity ratio measurements over time in the presence (red) and absence (gray) of 10 μM Inh-172 by flow cytometry. Cellular intensity ratios are plotted against the time and the fits for a single-exponential decay function are shown for reference. The fitted quenching half-life value increases 45% in the presence of inhibitor. In conjunction with the results in Fig. 6 showing that CFTR-specific modulators accelerate quenching, these results demonstrate that the observed cellular quenching reaction is rate-limited by CFTR conductance.

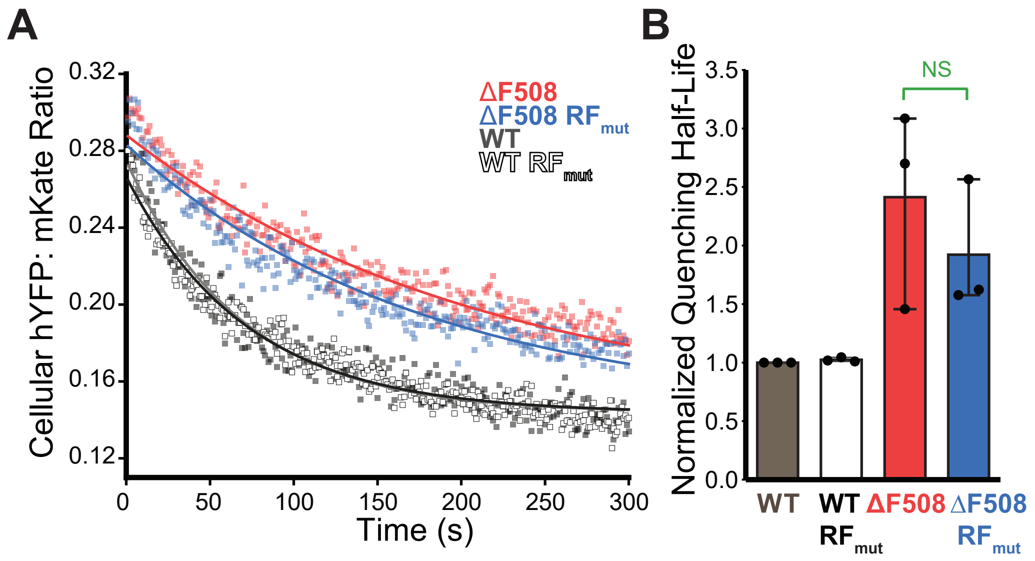


**Figure S11. CFTR Function in *EMC6* Knockout Cells.** The functional conductance of stably expressed CFTR variants was compared in the context of an *EMC6* knockout HEK293T cell line by measuring the time-dependent quenching of a halide-sensitive yellow fluorescent protein (hYFP). A) HEK293T cells stably expressing WT (white), WT $RF_{mut}$ (gray), ΔF508 (red), and ΔF508 $RF_{mut}$ (blue) were stimulated with 25 μM forskolin (0.06% DMSO Vehicle) to activate CFTR prior to measurement of the change in cellular hYFP: mKate intensity ratio measurements over time by flow cytometry. Cellular intensity ratios are plotted against the time and the global fits of the decay are shown for reference. B) A bar graph depicts the globally fit half-life for hYFP quenching for each variant. Values represent the average fitted values for each variant normalized relative to the WT value (n = 3). WT CFTR-mediated hYFP quenching is slower in *EMC6* knockout cells relative to parental HEK293T cells. As a result, the effect of the ΔF508 mutation appear less pronounced. Moreover, the $RF_{mut}$ modification has little, if any, impact on ΔF508 CFTR-mediated hYFP quenching in *EMC6* knockout cells- the *p*-value from a two-sample t-test indicated the difference between the half-lives derived from cells expressing these variants are not statistically significant.

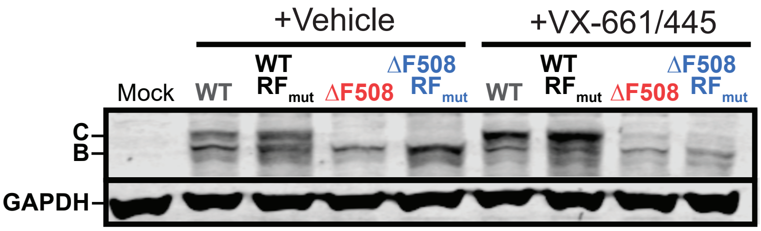


**Figure S12. Impact of ribosomal frameshifting on the pharmacological rescue of CFTR expression in HEK293T cells.** Western blotting was used to compare the expression and matu-ration of transiently expressed CFTR variants in the presence of CFTR modulators in HEK293T cells. A representative western blot depicting the relative abundance of the mature (band C) and immature (band B) CFTR glycoforms of each indicated variant in HEK293T cells treated for 16 hours with DMSO (Vehicle) or 3uM VX-661 + 3uM VX-445 is shown. A GAPDH loading control is included for reference. Though trends were consistent, quantitative variations in total CFTR expression levels in the presence of correctors were observed across western blots for different biological replicates. Trends were more precise when measured by flow cytometry due to the fact that this analysis is insensitive to variations in transfection efficiency. Nevertheless, C/B ratios from replicate experiements were quite consistent. For this reason, we chose to report expres-sion measurements from flow cytometry and maturation efficiencies from western blots in Fig. 6.

**Table S1. $Log_2$ (Fold-Change) in CFTR variant interactor identifications relative to WT.**

| Interactor | $WT\text{-}RF_{mut}$ | ΔF508 | $\Delta F508\text{-}RF_{mut}$ | Cluster ID |
|---|---|---|---|---|
| XPOT | 0.062 | 0.657 | 0.370 | 1 |
| EXOC5 | 0.245 | 0.765 | 0.556 | 1 |
| SAAL1 | 0.241 | 0.875 | 0.462 | 1 |
| EIF2B2 | -0.057 | 1.147 | 0.392 | 1 |
| UXS1 | -0.170 | 1.038 | 0.368 | 1 |
| PPP2R2A | 0.065 | 1.020 | 0.744 | 1 |
| SPTLC1 | 0.063 | 1.053 | 0.573 | 1 |
| IPO11 | 0.273 | -0.464 | 0.887 | 1 |
| HAX1 | 0.055 | 0.518 | 0.883 | 1 |
| FANCI | 0.143 | 0.142 | 0.561 | 1 |
| IGKC | -0.832 | 0.874 | -0.096 | 1 |
| IGKV3-20 | -0.602 | 1.015 | 0.041 | 1 |
| NCL | -0.237 | 1.353 | -0.643 | 1 |
| C4A | -0.403 | 1.045 | -0.568 | 1 |
| PSMC2 | 0.394 | 1.277 | 0.260 | 1 |
| RPL11 | 0.476 | 1.358 | 0.210 | 1 |
| PPP2R1A | 0.260 | 1.181 | 0.251 | 1 |
| DNAJB6 | 0.309 | 1.276 | 0.078 | 1 |
| RSL1D1 | 0.682 | 1.053 | 0.427 | 1 |
| PSMC1 | 0.280 | 1.548 | 0.424 | 1 |
| RPL24 | 0.593 | 1.431 | 0.653 | 1 |
| EFTUD2 | 0.150 | 1.060 | 0.057 | 1 |
| AHCY | -0.027 | 1.270 | 0.074 | 1 |
| HBA1 | -0.057 | 1.079 | 0.134 | 1 |
| RPS10 | 0.730 | 1.362 | -0.188 | 1 |
| CSNK2A1 | 0.305 | 0.958 | -0.168 | 1 |
| CPSF6 | 0.295 | 0.943 | -0.118 | 1 |
| MYCBP2 | 0.360 | 1.166 | -0.267 | 1 |
| RPL4 | 0.099 | 1.176 | -0.178 | 1 |
| ILKAP | 0.558 | -2.031 | -6.226 | 1 |
| SUN2 | 0.324 | 0.203 | -1.903 | 1 |
| AIFM1 | 0.289 | 0.205 | -1.465 | 1 |
| CCT5 | 0.365 | 0.264 | -1.673 | 1 |
| FLNA | 0.189 | 0.342 | -1.614 | 1 |
| PLCG1 | 0.509 | -0.454 | -3.406 | 1 |
| DDX20 | 0.284 | -0.103 | -2.230 | 1 |
| BBLN | -0.113 | -0.840 | -2.357 | 1 |
| CSE1L | 0.461 | -0.347 | -0.531 | 1 |
| IPO4 | 0.147 | -0.053 | -1.244 | 1 |
| KPNB1 | -0.084 | -0.501 | -0.994 | 1 |
| IPO7 | 0.060 | -0.225 | -0.870 | 1 |

| Interactor | WT-$RF_{mut}$ | ΔF508 | ΔF508-$RF_{mut}$ | Cluster ID |
|---|---|---|---|---|
| SNX4 | -0.012 | 0.166 | -0.389 | 1 |
| IPO9 | -0.090 | 0.051 | -0.571 | 1 |
| RPL36A | -0.114 | 0.054 | -0.452 | 1 |
| XPO1 | 0.304 | -0.002 | -0.429 | 1 |
| TNPO2 | 0.068 | 0.103 | -0.589 | 1 |
| RTN3 | 0.134 | -0.012 | -0.511 | 1 |
| CFTR | 0.000 | 0.000 | 0.000 | 1 |
| IPO8 | 0.099 | -0.166 | -0.077 | 1 |
| TNPO3 | 0.239 | 0.214 | 0.116 | 1 |
| TBC1D4 | 0.080 | 0.597 | -0.026 | 1 |
| GTF2I | 0.292 | 1.853 | -0.789 | 1 |
| RPL3 | 0.318 | 1.785 | -0.463 | 1 |
| FAF2 | 0.373 | 1.547 | -1.147 | 1 |
| SKP1 | 0.365 | 1.521 | -0.809 | 1 |
| RPL27 | 0.599 | 1.463 | -0.694 | 1 |
| RPS19 | 0.766 | 1.096 | -1.335 | 1 |
| LBR | 0.048 | 1.157 | -1.186 | 1 |
| PRPF6 | 0.406 | 0.472 | -1.202 | 1 |
| SYS1 | 0.353 | 0.389 | -0.852 | 1 |
| HSPA6 | 0.468 | 0.420 | -1.010 | 1 |
| SNRPB2 | 0.475 | 0.462 | -0.878 | 1 |
| THADA | 0.078 | 0.515 | -0.655 | 1 |
| UBE3C | 0.002 | 0.561 | -0.540 | 1 |
| TUBGCP2 | 0.389 | 1.057 | -0.722 | 1 |
| PUF60 | 0.348 | 0.651 | -0.813 | 1 |
| RPS24 | 0.423 | 0.697 | -0.674 | 1 |
| RARS1 | 0.363 | 0.785 | -0.580 | 1 |
| SLC25A6 | 0.371 | 0.681 | -0.539 | 1 |
| ESS2 | 0.380 | 1.615 | 2.707 | 2 |
| DARS1 | 0.286 | 1.536 | 2.743 | 2 |
| NDUFS3 | 0.236 | 1.508 | 2.695 | 2 |
| GNB1 | 0.191 | 1.532 | 2.660 | 2 |
| PSMD2 | 0.286 | 1.499 | 2.553 | 2 |
| SRSF3 | 0.350 | 1.504 | 2.525 | 2 |
| SF3A1 | 0.105 | 1.473 | 2.646 | 2 |
| TFIP11 | 0.257 | 1.438 | 2.585 | 2 |
| ENO1 | 0.195 | 1.457 | 2.581 | 2 |
| PGRMC1 | 0.284 | 1.150 | 2.731 | 2 |
| AUP1 | 0.482 | 1.281 | 2.644 | 2 |
| COPB1 | 0.188 | 1.275 | 2.495 | 2 |
| KCTD3 | 0.417 | 1.297 | 2.485 | 2 |
| SURF4 | 0.360 | 1.362 | 2.351 | 2 |
| APOA1 | -0.286 | 1.872 | 2.604 | 2 |

| Interactor | WT-RF$_{mut}$ | ΔF508 | ΔF508-RF$_{mut}$ | Cluster ID |
|---|---|---|---|---|
| CKMT1 | 0.047 | 1.409 | 2.514 | 2 |
| EIF3E | -0.043 | 1.470 | 2.457 | 2 |
| WRNIP1 | 0.023 | 1.441 | 2.388 | 2 |
| YWHAQ | 0.244 | 1.481 | 2.181 | 2 |
| XIAP | 0.254 | 1.459 | 2.297 | 2 |
| SNRPD1 | 0.280 | 1.443 | 2.301 | 2 |
| CDKN2A | 0.163 | 1.438 | 2.359 | 2 |
| WNK1 | 0.237 | 1.483 | 2.481 | 2 |
| ARHGEF2 | 0.170 | 1.492 | 2.429 | 2 |
| MTDH | 0.315 | 1.683 | 2.381 | 2 |
| XRN2 | 0.158 | 1.600 | 2.342 | 2 |
| MARS1 | 0.499 | 1.700 | 3.209 | 2 |
| MRPS27 | 0.495 | 1.816 | 2.877 | 2 |
| RSRC1 | 0.280 | 1.948 | 3.109 | 2 |
| CKAP4 | 0.320 | 1.423 | 2.924 | 2 |
| STAU1 | 0.240 | 1.468 | 3.054 | 2 |
| SRP9 | 0.408 | 1.557 | 3.059 | 2 |
| RBMX | 0.530 | 1.576 | 2.888 | 2 |
| DIAPH1 | 0.418 | 1.554 | 2.872 | 2 |
| DYNC1LI1 | 0.422 | 1.565 | 2.944 | 2 |
| PSMA1 | 0.346 | 1.815 | 3.568 | 2 |
| CLPX | 0.259 | 1.532 | 3.563 | 2 |
| NAP1L1 | 0.047 | 1.442 | 3.725 | 2 |
| RPL13A | 0.067 | 1.438 | 3.533 | 2 |
| OAT | 0.117 | 0.980 | 2.975 | 2 |
| NDUFA5 | 0.314 | 1.272 | 3.192 | 2 |
| KPNA2 | 0.188 | 1.411 | 3.266 | 2 |
| PSMD1 | 0.046 | 1.471 | 3.192 | 2 |
| TUBB1 | 0.049 | 1.479 | 1.089 | 3 |
| BAG2 | 0.106 | 1.766 | 1.244 | 3 |
| FLOT1 | 0.165 | 1.739 | 1.132 | 3 |
| ATP6V0D1 | 0.176 | 1.738 | 1.009 | 3 |
| ATAD3A | 0.358 | 2.076 | 0.381 | 3 |
| VCP | -0.109 | 1.901 | 0.838 | 3 |
| CANX | 0.158 | 1.816 | 0.751 | 3 |
| UBA52 | 0.085 | 1.570 | 0.648 | 3 |
| RCN2 | 0.303 | 1.133 | 1.204 | 3 |
| NUP93 | 0.260 | 1.076 | 1.279 | 3 |
| H1-4 | 0.584 | 1.298 | 1.075 | 3 |
| SDF4 | 0.440 | 0.954 | 0.835 | 3 |
| CLPB | 0.243 | 1.213 | 0.985 | 3 |
| SEC61A1 | 0.351 | 1.122 | 0.972 | 3 |
| TECR | 0.085 | 1.160 | 1.586 | 3 |

| Interactor | WT-RF$_{mut}$ | ΔF508 | ΔF508-RF$_{mut}$ | Cluster ID |
|---|---|---|---|---|
| ATP2A2 | 0.273 | 1.149 | 1.634 | 3 |
| FANCD2 | 0.280 | 1.189 | 1.597 | 3 |
| HEATR1 | -0.012 | 0.775 | 1.655 | 3 |
| SMC2 | 0.213 | 1.032 | 1.487 | 3 |
| TNPO1 | 0.209 | 0.952 | 1.500 | 3 |
| CIAO2B | 0.209 | 0.958 | 1.475 | 3 |
| HACD3 | 0.269 | 1.354 | 1.278 | 3 |
| HSD17B11 | 0.296 | 1.340 | 1.337 | 3 |
| PELO | 0.380 | 1.220 | 1.294 | 3 |
| MICOS13 | 0.383 | 1.317 | 1.330 | 3 |
| ATP1A1 | 0.311 | 1.325 | 1.419 | 3 |
| CHD4 | 0.328 | 1.266 | 1.442 | 3 |
| COPG1 | 0.251 | 1.352 | 1.509 | 3 |
| RAB10 | 0.192 | 1.351 | 1.475 | 3 |
| POLG | -0.209 | 1.579 | 1.410 | 3 |
| PSMC4 | 0.234 | 1.588 | 1.374 | 3 |
| RPS6KA2 | 0.108 | 1.471 | 1.388 | 3 |
| PCNA | 0.130 | 1.256 | 1.273 | 3 |
| DDX17 | 0.110 | 1.304 | 1.387 | 3 |
| MCCC2 | 0.028 | 1.229 | 1.359 | 3 |
| ERLIN1 | -0.046 | 1.332 | 1.571 | 3 |
| HELLS | -0.092 | 1.307 | 1.600 | 3 |
| AGK | -0.004 | 1.392 | 1.495 | 3 |
| HNRNPK | 0.092 | 1.280 | 1.469 | 3 |
| CNN3 | 0.033 | 1.314 | 1.479 | 3 |
| IARS1 | 0.393 | 1.599 | 1.586 | 3 |
| CYCS | 0.337 | 1.533 | 1.484 | 3 |
| EXOSC6 | 0.543 | 1.481 | 1.387 | 3 |
| TCOF1 | 0.563 | 1.493 | 1.594 | 3 |
| AP2B1 | 0.485 | 1.560 | 1.497 | 3 |
| FLG | 0.452 | 1.471 | 1.535 | 3 |
| MAP4K4 | 0.315 | 2.105 | 1.338 | 3 |
| FLOT2 | 0.451 | 1.889 | 1.705 | 3 |
| RPL32 | 0.394 | 1.770 | 1.552 | 3 |
| WDR83 | 0.459 | 1.770 | 1.516 | 3 |
| SNW1 | 0.363 | 1.437 | 1.638 | 3 |
| ACADM | 0.418 | 1.300 | 1.685 | 3 |
| SLC25A11 | 0.374 | 1.329 | 1.628 | 3 |
| USP9X | 0.317 | 1.331 | 1.743 | 3 |
| PYCR1 | 0.303 | 1.241 | 1.729 | 3 |
| VPS35 | 0.258 | 1.373 | 1.674 | 3 |

| Interactor | WT-RF$_{mut}$ | ΔF508 | ΔF508-RF$_{mut}$ | Cluster ID |
|---|---|---|---|---|
| EIF3C | 0.236 | 1.387 | 1.742 | 3 |
| TRMT112 | 0.270 | 1.358 | 1.792 | 3 |
| SMARCC2 | 0.209 | 1.655 | 1.676 | 3 |
| PSMC5 | 0.201 | 1.471 | 1.599 | 3 |
| ERP44 | 0.188 | 1.524 | 1.550 | 3 |
| LUC7L3 | 0.166 | 1.474 | 1.684 | 3 |
| PSMD3 | 0.234 | 1.550 | 1.639 | 3 |
| NUDT21 | 0.211 | 1.516 | 1.680 | 3 |
| PRMT5 | 0.187 | 1.513 | 1.642 | 3 |
| DDX21 | 0.184 | 1.277 | 1.763 | 3 |
| DHX15 | 0.072 | 1.337 | 1.785 | 3 |
| MMTAG2 | 0.056 | 1.239 | 1.753 | 3 |
| RBM25 | 0.055 | 1.469 | 1.716 | 3 |
| PSMC6 | 0.120 | 1.357 | 1.592 | 3 |
| HSDL2 | 0.153 | 1.352 | 1.637 | 3 |
| LYAR | 0.052 | 1.376 | 1.607 | 3 |
| PPP6R3 | 0.029 | 1.290 | 1.629 | 3 |
| SNRNP200 | 1.589 | 2.252 | 2.040 | 3 |
| TUBA4A | 1.737 | 2.141 | 1.894 | 3 |
| PPFIA3 | 0.270 | 2.837 | 1.897 | 3 |
| PTMA | 0.759 | 2.774 | 1.649 | 3 |
| GGCT | 0.946 | 1.699 | 1.901 | 3 |
| CEP170 | 0.471 | 1.744 | 1.922 | 3 |
| SPAG9 | 0.534 | 1.631 | 1.990 | 3 |
| EEF1B2 | 0.703 | 1.529 | 1.827 | 3 |
| SRP14 | 0.569 | 1.526 | 1.936 | 3 |
| SRSF1 | 0.489 | 1.449 | 1.954 | 3 |
| ATP5PD | 0.608 | 1.423 | 1.819 | 3 |
| CLPTM1 | 0.507 | 1.371 | 1.840 | 3 |
| HSPE1 | 0.582 | 2.162 | 2.232 | 3 |
| NUDT16L1 | 0.536 | 1.594 | 2.489 | 3 |
| NUDC | 0.848 | 1.641 | 2.197 | 3 |
| NDUFA8 | 0.599 | 1.582 | 2.132 | 3 |
| HSPB1 | 0.656 | 1.745 | 2.180 | 3 |
| C6orf47 | 0.613 | 1.673 | 2.334 | 3 |
| ALDOA | 0.329 | 1.684 | 2.069 | 3 |
| CWC25 | 0.350 | 1.561 | 2.131 | 3 |
| RSRC2 | 0.279 | 1.598 | 2.104 | 3 |
| PSMA3 | 0.212 | 1.567 | 2.086 | 3 |
| RBBP1 | 0.293 | 1.497 | 1.722 | 3 |
| LRPAP1 | 0.309 | 1.481 | 1.780 | 3 |

| Interactor | WT-$RF_{mut}$ | ΔF508 | ΔF508-$RF_{mut}$ | Cluster ID |
|---|---|---|---|---|
| DYNC1LI2 | 0.275 | 1.450 | 1.771 | 3 |
| SYNCRIP | 0.323 | 1.520 | 1.883 | 3 |
| PRDX6 | 0.395 | 1.544 | 1.883 | 3 |
| OGFR | 0.392 | 1.497 | 1.805 | 3 |
| RAB11A | 0.389 | 1.417 | 1.791 | 3 |
| ERLIN2 | 0.162 | 1.633 | 1.935 | 3 |
| MARK2 | 0.178 | 1.709 | 1.838 | 3 |
| PRPF19 | 0.223 | 1.483 | 1.866 | 3 |
| SPTLC2 | 0.198 | 1.454 | 1.910 | 3 |
| PRPF40A | 0.213 | 1.501 | 1.795 | 3 |
| SMARCD1 | 0.243 | 1.568 | 1.868 | 3 |
| RPN1 | 0.365 | 1.423 | 2.001 | 3 |
| HTATSF1 | 0.248 | 1.498 | 1.937 | 3 |
| SLC25A1 | 0.297 | 1.482 | 1.976 | 3 |
| SLC16A1 | 0.453 | 1.087 | 1.883 | 3 |
| SNRPB | 0.428 | 1.372 | 2.163 | 3 |
| ARHGEF1 | 0.402 | 1.292 | 2.131 | 3 |
| ABCD3 | 0.248 | 1.180 | 2.161 | 3 |
| MAGED1 | 0.393 | 1.069 | 2.213 | 3 |
| LYZ | -0.219 | 1.004 | 2.163 | 3 |
| DCXR | 0.234 | 1.400 | 1.984 | 3 |
| DNAJA3 | 0.207 | 1.305 | 2.070 | 3 |
| CAD | 0.182 | 1.373 | 2.071 | 3 |
| CNOT1 | 0.243 | 1.398 | 2.068 | 3 |
| RPL15 | 0.084 | 1.471 | 2.211 | 3 |
| TIM23B | 0.035 | 1.429 | 2.173 | 3 |
| ALDH18A1 | 0.105 | 1.363 | 2.021 | 3 |
| TNRC6B | 0.085 | 1.400 | 2.077 | 3 |
| HNRNPM | 0.087 | 1.414 | 2.105 | 3 |
| FN | 0.041 | 1.411 | 1.942 | 3 |
| HSPA4 | 0.134 | 1.511 | 1.997 | 3 |
| SF3A3 | 0.089 | 1.505 | 2.039 | 3 |
